# Discovery of an evolutionarily conserved smooth muscle cell-specific lncRNA *CARMN*

**DOI:** 10.1101/2020.01.30.927335

**Authors:** Kunzhe Dong, Jian Shen, Xiangqin He, Liang Wang, Guoqing Hu, Kirstopher M. Bunting, Rachael Dixon-Melvin, Zeqi Zheng, Hongbo Xin, Meixiang Xiang, Almira Vazdarjanova, Jiliang Zhou

**Affiliations:** Department of Pharmacology and Toxicology, Medical College of Georgia, Augusta University, Augusta, GA 30912; Department of Cardiology, The Second Affiliated Hospital, Zhejiang University, Hangzhou, Zhejiang, 310009, China; The National Engineering Research Center for Bioengineering Drugs and the Technologies, Institute of Translational Medicine, Nanchang University, Nanchang, Jiangxi, 330031, China; School of Life Sciences, Nanchang University, Nanchang, Jiangxi, 330031, China; Department of Cardiology, The First Affiliated Hospital of Nanchang University, Nanchang, Jiangxi, 330006, China

## Abstract

Differentiated vascular smooth muscle cells (VSMCs) are critical in maintaining vascular homeostasis by expressing a unique repertoire of contractile genes. Despite the well-defined coding transcriptome in differentiated VSMCs, little is known about the non-coding gene expression signature. Herein, by de novo analyzing publicly available RNA-seq and single cell RNA-seq datasets generated from different tissues and cell types, we unambiguously identified *CARMN* (CARdiac Mesoderm Enhancer-associated Non-coding RNA) as an evolutionarily conserved, SMC-specific lncRNA. *CARMN* was initially annotated as the host gene of *MIR143/145* cluster and recently reported to play roles in cardiac differentiation. Here, we generated a *Carmn* GFP knock-in reporter mouse model and confirmed its specific expression in SMCs *in vivo*. In addition, we found *Carmn* is transcribed independently from *Mir143/145* and only expressed transiently in embryonic cardiomyocytes and thereafter becomes restricted to adult SMCs in both human and mouse. Furthermore, we demonstrated that *CARMN* expression is not only dramatically decreased in human vascular diseases but functionally critical in maintaining VSMC contractile phenotype *in vitro*. In conclusion, we provided the first evidence showing that *CARMN* is an evolutionarily conserved SMC-specific lncRNA, down-regulated in different human vascular diseases, and a key lncRNA for maintaining SMC contractile phenotype.

## Introduction

Smooth muscle cells (SMCs) are the major contractile component of most hollow organs, such as blood vessels, bladder, colon and esophagus. Fully differentiated SMCs are characterized by the presence of a unique repertoire of contractile proteins including CNN1 (1), TGFB1I1 (2), TAGLN (3), LMOD1 (4), ACTA2 (5), 130-kDa myosin light chain kinase (MYLK) (6) and smooth muscle myosin heavy chain (MYH11) (7). In many vascular diseases such as atherosclerosis and restenosis, vascular SMCs (VSMCs) switch from a contractile to a synthetic phenotype that is characterized by increased motility and proliferation, as well as reduced expression of these SMC-specific contractile proteins (8). While expression of these SM-specific genes in normal and diseased VSMCs has been well characterized, little is known about SM specifically expressed long non-coding RNAs (lncRNAs).

LncRNAs, defined as transcripts larger than 200 nucleotides and with no apparent protein-coding potential, have been found to be pervasively transcribed in >80% of human genome, outstripping the <3% of protein-coding genomic DNA (9). Although initially regarded as “junk”, accumulating evidence suggests that many of them play critical roles under variety of pathological conditions (9). Unlike protein-coding genes that are well-studied in SM biology, our understanding of the expression and function of lncRNAs in SMCs remains in its infancy (10). In addition to several lncRNAs which have been implicated in SM biology (11, 12), our recent study demonstrated that the lncRNA *NEAT1* plays a critical role in smooth muscle de-differentiation (13). However, many of these lncRNAs are not conserved between species, and not exclusively expressed in SMCs and importantly, their expression has not been carefully tracked by *in vivo* reporter systems. These limitations collectively hinder our understanding of the importance of lncRNAs in SMCs.

The technical advances in whole transcriptome RNA-seq (bulk RNA-seq), and especially single cell RNA-seq (scRNA-seq), have offered an unprecedented opportunity for discovering tissue/cell-specific genes in high-throughput and unbiased manners (14). In this study, we used an *in silico* approach to probe unbiased proprietary and diverse publicly available bulk RNA-seq and scRNA-seq datasets to search for SMC-specific lncRNAs. This search identified *CARMN* (CARdiac Mesoderm Enhancer-associated Non-coding RNA) as an evolutionarily conserved SMC-specific lncRNA. *CARMN* was recently reported to play roles in cardiac differentiation (15, 16), and was initially annotated as a lncRNA expressed from the *MIR143/145* cluster locus, the best characterized microRNAs in regulating SM differentiation and phenotypic modulation (12, 17-20). We generated a GFP knock-in (KI) mouse model which expresses GFP under the control of the endogenous *Carmn* gene promoter in order to visualize *Carmn* expression *in vivo.* Analysis of these mice confirmed *Carmn* expression specifically in SMCs. In addition, we provide evidence that *Carmn* is transcribed independently with *Mir143/145* cluster. Furthermore, we found *CARMN* is expressed transiently in cardiomyocytes during early embryonic development and thereafter becomes restricted to postnatal SMCs in both human and mouse. Importantly, we found that *CARMN* expression is dramatically decreased in various human vascular diseases such as atherosclerosis and aneurysm as well as rodent vascular disease models. We further demonstrated that the knock down of endogenous *CARMN* in human coronary artery SMCs (HCASMCs) significantly promotes SMC proliferation and migration, and inhibits the expression of SMC contractile proteins, while over-expression of *Carmn* had opposite effects on SMCs. These findings collectively suggest that *CARMN* is a key regulator of SMC phenotype and represents a promising therapeutic target for treatment of SMC-related diseases.

## Results and Discussion

To identify SMC-enriched lncRNAs, we analyzed existing public bulk RNA-seq datasets generated from human VSMCs and 5 different non-SMC cell types. From this unbiased analysis we identified the top 15 lncRNAs that are most enriched in VSMCs (**Figure 1A**). Analysis of another independent bulk RNA-seq dataset for 22 different human primary cell types from the ENCODE project identified 4 lncRNAs preferentially expressed in VSMCs (**Figure 1B** and **Supplemental Table I**). Interestingly, *CARMN* was the only lncRNA that was identified within both datasets. Furthermore, ChIP-seq data from the ENCODE database revealed that, in contrast to human vein endothelial cells (HUVECs), transcriptional activity in the *CARMN* gene locus is high in human aortic VSMCs as indicated by the presence of high levels of H3K27ac (an epigenetic mark of active chromatin) and undetectable levels of H3K27me3 (an epigenetic mark of repressed chromatin). In contrast, high levels of H3K27me3 were present at this locus in HUVECs (**Figure 1C**). Interestingly, examination of two additional histone modifications of *CARMN* locus in SMCs, H3K4me2 and H3K4me3, which are known to be enriched at the transcription start site and commonly used as markers for active gene promoter activity (21), identified a potentially independent promoter upstream of the *MIR143/145* which was distinct from the active promoter signature upstream of the *CARMN* locus (**Supplemental Figure 1A**). This observation suggests that, although a human *CARMN* transcript harboring *MIR143/145* is annotated in the UCSC genome database, *MIR143/145* may be transcribed independent of *CARMN*, a common scenario observed in one-third of the miRNAs located within protein-coding or non-coding genes (22). These findings are also in accord with the recent studies showing that *CARMN* functions in a manner independent of *MIR143/145* in regulating human cardiac specification and differentiation (15, 16). Further analysis of GTEx database revealed that *CARMN* is mostly expressed in vascular tissues such as thoracic aorta, coronary and tibial artery, followed by SM-enriched gastrointestinal (GI) tissues but not in the heart in adult humans (**Figure 1D**). We next de novo analyzed a scRNA-seq dataset that was generated from coronary artery of four human cardiac transplant recipients (23). Visualization of the expression of *CARMN* at the single-cell level revealed that *CARMN* expression is restricted to SMCs and pericytes, a type of cell sharing expression of many SMC markers (24). The restricted expression pattern of *CARMN* is in parallel with well-known SMC markers including *MYH11* (**Figure 1E**), *ACTA2*, *CNN1* and *TAGLN* (**Supplemental Figure 1B**), confirming the specific expression of *CARMN* in SMCs. This analysis also revealed that *CARMN* expression was virtually down-regulated in phenotypically modulated SMCs compared to the normal SMCs (**Figure 1E**). Taken together, these unbiased large-scale bioinformatic analysis demonstrate *CARMN* is a SMC- and SM tissue-enriched lncRNA in humans.

**Figure 1.**
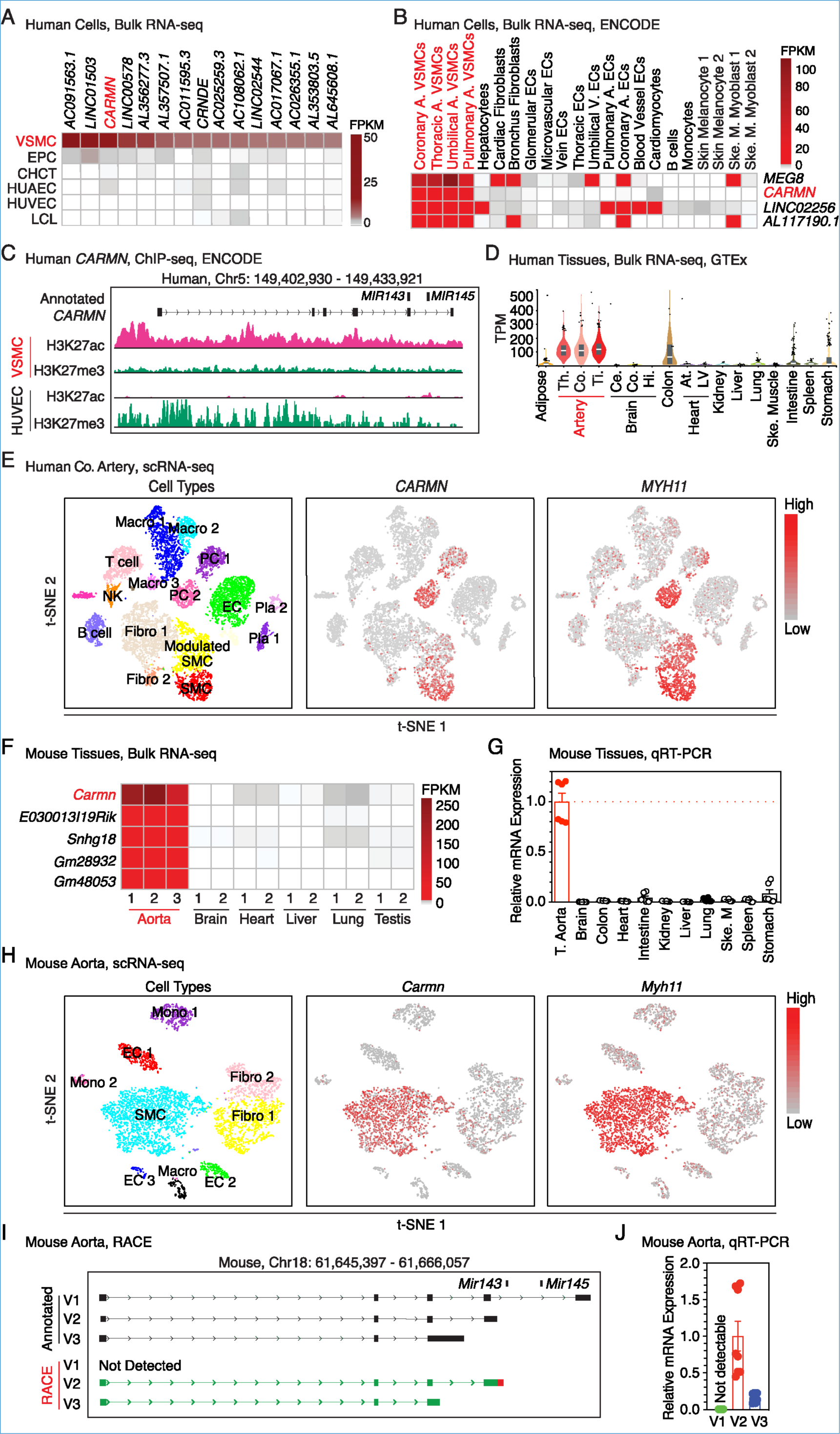
Identification of *CARMN* as an evolutionarily conserved SMC-specific lncRNA. (**A**) Heatmap showing 15 VSMC-enriched lncRNAs identified in the publicly available human RNA-seq dataset. EPC: endothelial progenitor cells; CHCT: chondrocytes; HUAEC: human aortic endothelial cells; HUVEC: human umbilical vein endothelial cells; LCL: lymphoblastoid cell lines. FPKM: Fragments Per Kilobase of transcript per Million mapped reads. (**B**) Heatmap showing 4 VSMC-enriched lncRNAs in 22 primary human cell types retrieved from the ENCODE database. A: artery; V: Vein; ECs: endothelial cells; Ske. M.: Skeletal Muscle. (**C**) ChIP-seq tracks of histone 3 modifications at the human *CARMN* locus in aortic VSMC vs HUVEC as revealed by the ENCODE project. Y-axis scale: 20 for H3K27ac and 5 for H3K27me3. (**D**) *CARMN* expression in human tissues was revealed by the GTEx database. Th: thoracic; Co: coronary; Ti: tibial artery; Ce: cerebellum; Co: cortex; Hi.: hippocampus; At: atrium; LV: left ventricle. TPM: Transcripts Per Kilobase Million. (**E**) t-SNE (t-distributed stochastic neighbor embedding) visualization of cell types isolated from the right coronary artery of four human patients (left), and the expression of *CARMN* (middle) and *MYH11* (right) revealed by the scRNA-seq analysis. The color scale on the right indicates the expression levels of both *CARMN* and *MYHH11.* Fibro: fibroblast; EC: endothelial cell; PC: pericyte cell; Macro: macrophage; NK: natural killer cell; Pla: plasma cell. (**F**) Heatmap showing the aorta-enriched lncRNAs in mouse identified by RNA-seq data of different mouse tissues. (**G**) *Carmn* expression was assessed in different mouse tissues by qRT-PCR. The expression of *Carmn* in mouse thoracic aorta was set to 1. N=6. T, thoracic aorta; Ske. M: skeletal muscle. (**H**) t-SNE visualization of cell types present in normal mouse aorta (left) and the expression of *Carmn* (middle) and *Myh11* (right) revealed by scRNA-seq. The color scale on the right indicates the expression levels of both *Carmn* and *Myh11.* Macro: macrophage. (**I**) *Carmn* transcripts were identified by RACE and (**J**) qRT-PCR validated that V2 is the most abundant transcript in mouse aorta. N=6.

To identify lncRNAs preferentially expressed in mouse aorta, we next re-analyzed bulk RNA-seq generated from adult mouse aortic tissues and 5 other non-SM tissues in mouse. This analysis revealed that *Carmn* is the most abundant aorta-enriched lncRNA in mouse (**Figure 1F**). Similar to the findings in humans, qRT-PCR validated *Carmn* is most abundantly expressed in thoracic aorta, followed by SM-enriched GI tissues such as intestine and stomach but not in the heart of adult mice (**Figure 1G**). To further visualize *Carmn* expression with single-cell resolution, we de novo analyzed the published scRNA-seq data generated from normal adult mouse aorta. This analysis showed that similar to well-known SM contractile markers including *Myh11* (**Figure 1H**), *Acta2*, *Cnn1* and *Tagln*, *Carmn* is selectively expressed in aortic SMCs (**Supplemental Figure 1C**). This finding is consistent with the observations in human coronary artery (**Figure 1E**). As an initial step to explore *Carmn* transcription in SMCs, we observed that three splice variants of mouse *Carmn* were currently annotated in UCSC genome database, with transcript V1 encompassing *Mir143* and *Mir145* (**Figure 1I**). However, our RACE (Rapid Amplification of cDNA Ends) assay using adult mouse aorta cDNA was unable to detect annotated V1 but confirmed the presence of the V2 and V3 transcripts. Our V2 (1228 bp) transcript had an extended 3’ untranslated region (UTR) (GenBank accession number: MN904529) relative to the previously annotated V2 transcript, and V3 (759 bp) was shorter than the previously annotated transcript (GenBank accession number: MN904530). Sequencing data of these amended transcripts revealed that neither of them contain any sequence of *Mir143* or *Mir145* (**Figure 1I**), indicating that *Carmn* is not the precursor transcript of *Mir143/145* cluster and is independently transcribed, at least in mouse aorta. To further confirm this observation, we examined the alternative splicing events of mouse *Carmn* gene revealed by bulk RNA-seq of mouse aorta. The analysis revealed that splicing junctions cover our amended transcripts V2 and V3, as well as its neighboring gene *Bvht*, while no junction reads span exon 4 and exon 5 that is unique to the annotated transcript V1 (**Supplemental Figure 1D**). This unbiased *in silico* analysis supported our RACE results and further demonstrated that annotated *Carmn* transcript V1 encompassing *Mir143/145* does not exist in adult mouse aorta. Further qRT-PCR data revealed that the novel V2 transcript is the most abundant one in mouse aorta, followed by V3 while the annotated V1 is undetectable (**Figure 1J**). Although there is no *Carmn* gene annotated in current rat genome, a portion of rat *Carmn* cDNA sequence that is positionally conserved *to Mir143/145*, was successfully cloned and confirmed by Sanger sequencing using primers located within the homologous region of mouse and rat *Carmn* (**Supplemental Figure 1E-F**). Taken together, by combining unbiased *in silico* analysis, we discovered that *Carmn* is an evolutionarily conserved SMC-specific lncRNA transcribed independently of *Mir143/145* cluster.

To visualize the SMC-specific expression of *Carmn in vivo*, we generated a *Carmn* GFP KI mouse model with a promoterless, reversed splicing acceptor (SA)-membrane bound GFP gene trap cassette, which is flanked by two pairs of oppositely orientated *Lox2272* sites (25) and *LoxP* sites (referred to as conditional, PFG, P mice). This KI cassette was placed in the intron between exon 2 and 3 of mouse *Carmn* gene (**Figure 2A**). Cre-mediated recombination will lead to inversion of the reversed GFP cassette (PFG to GFP) and deletion of *Carmn* exon 2. As a result, *Carmn* transcription will be prematurely terminated by splicing of exon 1 to the splice acceptor of the GFP cassette resulting in GFP expression driven by the endogenous *Carmn* gene promoter (referred to as GFP, G mice). GFP expression should therefore faithfully display the endogenous *Carmn* expression *in vivo.* To trace *Carmn* expression *in vivo*, we crossed the female *Carmn*^PFG/WT^ (PW) mice with male mice ubiquitously expressing *CMV*-Cre to invert the GFP cassette and delete *Carmn* exon 2 in all mouse tissues (**Figure 2B**). Genotyping results using tail DNA extracted from the offspring of *CMV*-Cre^+^/*Carmn*^GFP/WT^ (GW), *CMV*-Cre^-^/*Carmn*^PFG/WT^ (PW) and *CMV*-Cre^+^/*Carmn*^WT/WT^ (WW) mice confirmed Cre-mediated recombination within the *Carmn* gene locus in the GW mouse (**Figure 2C**). Although ubiquitous Cre-mediated recombination is a global event, data from Western blotting revealed that GFP expression is restricted to the aorta, while absent in heart, skeletal muscle and brain, in GW heterozygous mice (**Figure 2D**), confirming the preferential expression of *Carmn* in aortic tissue as revealed by the *in silico* analysis. Consistent with the Western blotting results, direct visualization of GFP fluorescence in adult mouse revealed specific expression in aorta and coronary arteries (**Figure 2E**), as well as in other hollow organs such as bladder, gallbladder, colon and intestine, and in regions of vascular SMCs in brain and quadriceps muscles, and bronchus in lung, in GW heterozygous mice (**Supplemental Figure 2A-H**). The preferential expression of *Carmn* in SMCs, but not in cardiomyocytes or skeletal muscle, was further confirmed by direct visualization of GFP in sections of adult GW mouse heart and skeletal muscle (**Supplemental Figure 2I**). No GFP signal was detected in any tissues in WW (WT) or PW (Cre negative) mice. To reveal the cell types labeled by GFP in aorta, we sectioned thoracic aorta of GW mouse for immunostaining of GFP, the SM-specific marker ACTA2, and the endothelial marker PECAM1. Data from immunostaining assays demonstrated that GFP, specifically expressed in the media layer of aorta, is colocalized with ACTA2, but not in endothelial or adventitial cells (**Figure 2F**). Data from RNA FISH (Fluorescent *in situ* hybridization) assays further revealed that *Carmn* expression is exclusively confined to the nuclei of SMCs in aorta, while is absent in endothelial cells and adventitial cells (**Figure 2G**). Similarly, *Carmn* is localized in the nuclei of cultured mouse VSMCs as revealed by RNA FISH assay (**Figure 2H**). The subcellular localization in the nuclei of SMCs is consistent with a proposed role for *Carmn* in regulating gene expression and/or nuclear organization (26). Collectively, these results provide direct *in vivo* evidence showing that *Carmn* is specifically expressed in the nuclei of SMCs.

**Figure 2.**
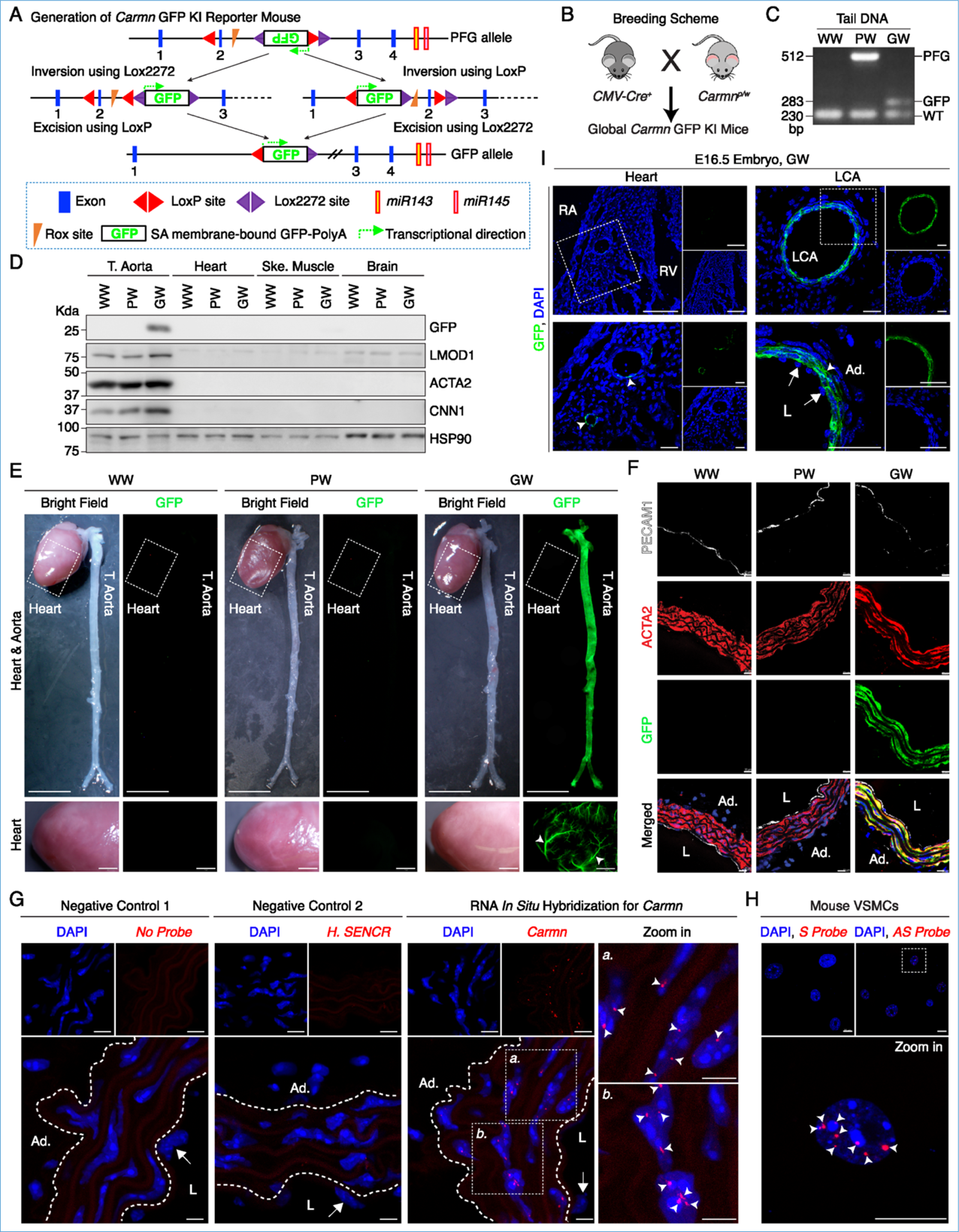
Visualization of SMC-specific expression of *Carmn in vivo*. (**A**) Strategy used to generate the *Carmn* GFP KI mouse model showing the inverted GFP KI cassette in the *Carmn* gene locus. SA: splicing acceptor site. (**B**) The breeding strategy to generate global *Carmn* GFP KI mice. (**C**) PCR analysis of genomic DNA showing recombination of the *Carmn* gene in DNA extracted from WW, PW and GW mice. (**D**) Western blot showing GFP expression specifically in thoracic aorta (T) of GW mice. Ske: skeletal muscle. (**E**) Heart and aorta were dissected from adult WW, PW and GW mice and photographed under bright field or GFP channel. T: thoracic aorta. Scale bar: 5 mm and 1 mm in upper and bottom panel, respectively. The arrowheads point to the representative coronary arteries in the heart of GW mouse. (**F**) Immuno-staining was performed to examine the specific cellular localization of GFP in thoracic aorta dissected from WW, PW and GW mice. L: lumen. Ad: adventitial layer. Scale bar: 10 μm. (**G**) RNA-FISH was carried out to visualize *Carmn* (right, red) in adult aorta. The sections hybridized with hybridization buffer only (left), human antisense (AS) *SENCR* probe (middle) were used as negative controls. Cell nuclei were counter stained with DAPI (blue). Dashed lines indicate external or internal elastic lamina. The boxed areas are magnified to the right. The arrows point to endothelial cells and the arrowheads indicate *Carmn* signal, respectively. L: lumen; Ad: adventitial layer. Scale bar: 10 μm. (**H**) *Carmn* subcellular localization in cultured mouse VSMCs was revealed by RNA-FISH using *Carmn* AS probe (red). *Carmn* sense probe (S) was used as negative control. Cell nuclei were stained with DAPI (blue). The boxed area is magnified below. The arrowheads indicate *Carmn* signal. Scale bar: 20 μm. (**I**) Visualization of GFP expression in E16.5 GW embryos showing that GFP is specifically expressed in arteriole SMCs/pericytes in right ventricle (RV) and medial SMCs (arrowheads) in left common carotid artery (LCA) but not in cardiomyocytes, endothelium (arrows) or adventitial (Ad.) cells. RA: right atrium; L: lumen; Ad: adventitial layer. Scale bar: 20 μm.

While our *in silico* analysis and KI reporter mouse model presented above demonstrate that *Carmn* expression is SMC-specific, two recent studies showed that *CARMN* plays roles in cardiac cell specification (15) and adult cardiac precursor cells *in vitro* (16). To test whether *CARMN* is expressed in cardiomyocytes during mouse embryonic development, we examined *Carmn* expression in GW embryos at embryonic day 16.5 (E16.5) and we found GFP is selectively expressed in the medial SM layers of carotid artery or in the arteriole SMCs or pericytes in the heart, not in endothelium or cardiomyocyte at this developmental stage (**Figure 2I**). Furthermore, we re-analyzed scRNA-seq dataset generated from cells isolated from mouse embryonic heart at E10.5, a time point in which many SMC-restricted genes are expressed transiently in the heart. Data from this de novo analysis showed that in mouse heart at E10.5, a portion of cardiomyocytes, as well as a cluster of cells expressing both cardiomyocyte and endothelial markers (Cluster 6), exhibit some level of *Carmn* expression (**Supplemental Figure 3A**). Analysis of scRNA-seq data of cardiomyocytes and non-cardiomyocytes isolated from adult mouse heart revealed that *Carmn* expression is absent in cardiomyocytes but restricted to SMCs/pericytes in adult mouse heart (**Supplemental Figure 3B-C**). Similarly, *CARMN* expression is observed at some extent in some cardiomyocytes, fibroblasts and endothelial cells in human developmental 6.5-7 post conception weeks (PCW) heart (**Supplemental Figure 3D**), while confined to SMCs and pericytes in adult human heart (**Supplemental Figure 4E**). These results collectively suggest that *CARMN* is transiently expressed in cardiomyocytes and other cell types during early development but becomes confined to SMCs as development progresses in both mouse and human. As our KI reporter mouse model has shown that at E16.5, *Carmn* expression displays SMC-specific but undetectable in cardiomyocytes (**Figure 2I**), it is likely that *Carmn* expression is inactivated in cardiomyocytes and becomes restricted to SMCs between E10.5 and E16.5.

Given the highly specific expression of *CARMN* in VSMCs and the observation that *CARMN* expression is reduced in modulated SMCs compared to contractile SMCs as revealed by the scRNA-seq of diseased human artery (**Figure 1E**), we next sought to further explore its relevance to vascular diseases, which exhibit SMC phenotypic switching from a contractile to a synthetic phenotype (8). Based on scRNA-seq data, the percentage of cells with detectable *CARMN* levels decreases from 62.5% in normal SMCs to 16.6% in modulated SMCs in diseased human coronary artery (**Figure 3A**). This is consistent with the dramatic down-regulation of *CARMN* in modulated SMCs (**Figure 3B**). Targeted re-analysis of previously published microarray or bulk RNA-seq datasets revealed that *CARMN* is significantly down-regulated in human atherosclerotic arteries and cerebral aneurysms, respectively (**Figure 3 C-D**). To extend these human findings to rodent vascular disease models, we re-analyzed scRNA-seq data of aortic roots and ascending aorta from *ApoE*^-/-^ mice at baseline before high-fat diet (HFD), as well as after 8 and 16 weeks of HFD feeding, which is a commonly used atherosclerotic mouse model. Our analysis revealed a marked reduction of percentage of *Carmn* positive cells as well as *Carmn* expression in modulated SMCs compared to normal SMCs during the disease progression (**Figure 3E**), similar to other SM-contractile markers including *Myh11*, *Acta2* and *Cnn1*. This expression pattern is opposite to SM de-differentiation markers such as *Lum* and *Tnfrsf11b* which are up-regulated in phenotypically modulated VSMCs (**Supplemental Figure 4A-C**). The down-regulation of *Carmn* in mouse atherosclerotic aortic tissues was independently validated by our qRT-PCR results (**Figure 3F**). Furthermore, qRT-PCR analysis revealed that *Carmn* expression is decreased in ligation-injured carotid arteries and wire-injured femoral arteries in mouse and balloon-injured carotid arteries in rat (**Figure 3G-I**). All of these rodent models mimic human vascular wall diseases involving VSMC phenotypic switching which drives neointima formation (27-29). Targeted re-analysis of previous RNA-seq studies revealed that *CARMN* was down-regulated in human saphenous vein SMCs following stimulation with PDGF-BB and IL-1*α*, two well-characterized chemokines that induce VSMC phenotypic switching (30, 31) (**Figure 3J**). Furthermore, data from qRT-PCR assays revealed that *CARMN* expression is down-regulated during de-differentiation of HCASMCs induced by switching the culture medium from a differentiation to a growth medium (**Figure 3K**). Re-analysis of RNA-seq of rat VSMCs revealed marked down-regulation of *Carmn* in rat VSMCs treated with PDGF-BB (**Figure 3L**), which was further validated by qRT-PCR analysis (**Figure 3M**). Collectively, these data suggest *CARMN* expression is decreased during VSMC phenotypic modulation in human diseased artery, in rodent models of vascular wall diseases and in cultured VSMCs in response to growth and inflammatory stimuli.

**Figure 3.**
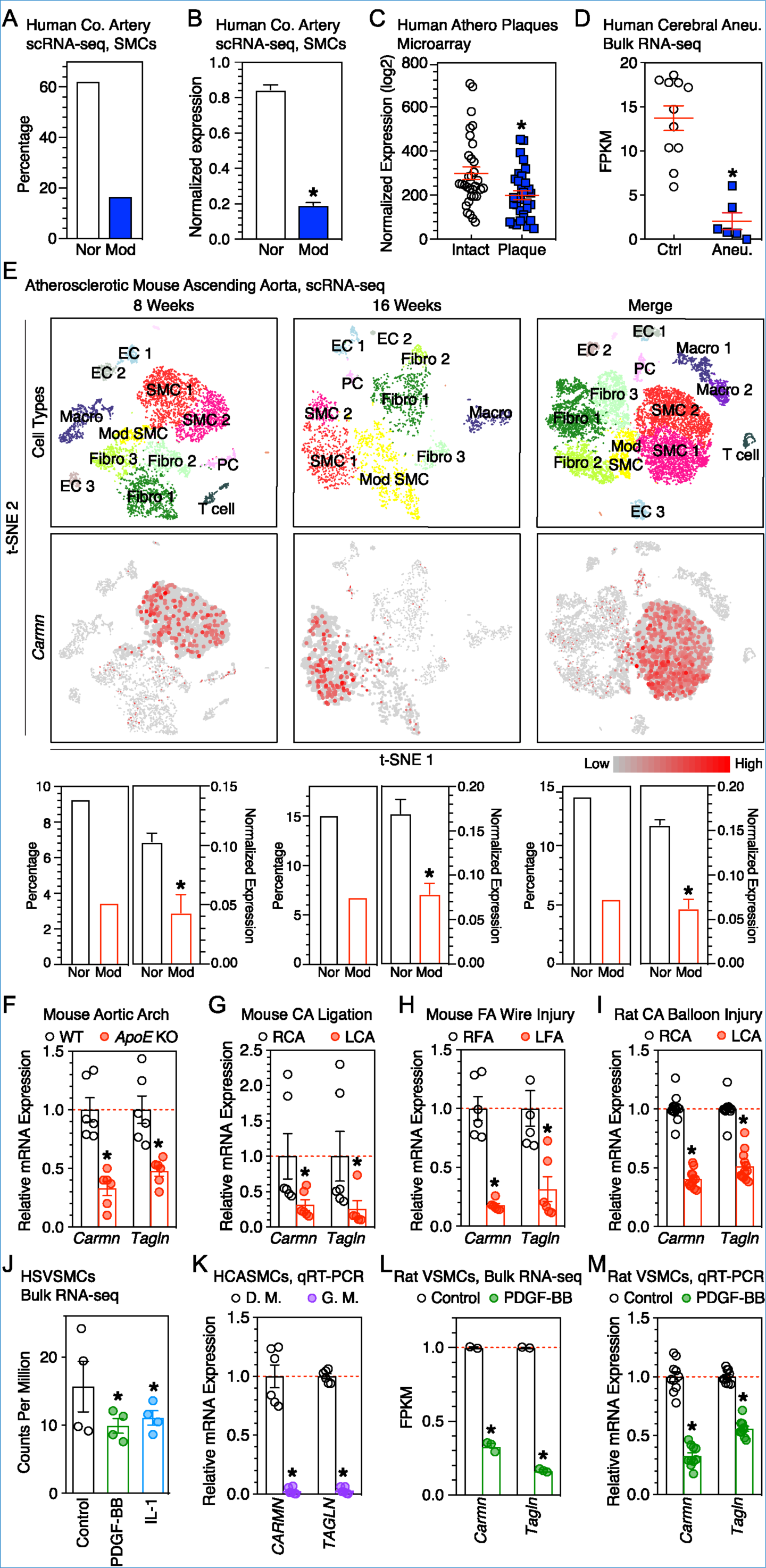
*CARMN* expression is down-regulated in human VSMC-related diseases, rodent vascular disease models and VSMCs in response to stimuli of phenotypic modulation. (**A**) Quantitative analysis of the percentage of *CARMN* positive cells (non-zero) and (**B**) expression level of *CARMN* in normal (Nor) and modulated (Mod) SMC clusters revealed by scRNA-seq of diseased human coronary (Co.) artery as shown in Figure 1E. *FDR (false discovery rate) < 0.05. (**C**) Re-analysis of previously reported microarray or bulk RNA-seq datasets to show *CARMN* expression significantly decreased in diseased human arteries such as atherosclerotic arteries and (**D**) cerebral aneurysm (Aneu.), respectively. Ctrl: control. *FDR < 0.05. (**E**) t-SNE visualization of cell types present in the mouse aortic root and ascending aorta after 8 (upper, left) and 16 weeks (upper, middle) of HFD, and merged cells from all time points including baseline before HFD, after 8 and 16 weeks of HFD feeding (upper, right), and the expression of *Carmn* (middle panel) at different time points as revealed by scRNA-seq. The bottom panel shows the quantitative analysis of percentage of *Carmn* positive cells (non-zero) and expression level of *Carmn* in normal (Nor) and modulated (Mod) SMC clusters. Fibro: fibroblast; EC: endothelial cell; PC: pericyte; Macro: macrophage. *FDR < 0.05. (**F**) qRT-PCR analysis of *Carmn* expression in lesion-laden aortic arch in *ApoE*^-/-^ mice following western-diet for 3 months or chow diet-fed control WT mice. *P < 0.05. (**G**) qRT-PCR analysis of *Carmn* expression in ligation-injured mouse left carotid artery (LCA), wire-injured mouse left femoral artery (LFA, **H**), balloon injured rat LCA (**I**). Uninjured right CA (RCA) or right FA (RFA) served as control (set to 1). N=6. *P < 0.05. (**J**) Re-analysis of public RNA-seq data showing decreased expression of *CARMN* upon stimulation of PDGF-BB or IL-1*α* in human saphenous vein (HSV) SMCs. *FDR < 0.05. (**K**) qRT-PCR to show *CARMN* expression in HCASMCs cultured with differentiation medium (D.M., set to 1) or growth medium (G.M.) for 48 hours. *P < 0.05. **(L)** Re-analysis of public RNA-seq data showing reduced expression of *Carmn* in rat VSMCs treated with PDGF-BB for 24 hours. *FDR < 0.05. (**M**) qRT-PCR analysis of *Carmn* expression in rat VSMCs treated with PDGF-BB (50ng/ml) for 24 hours. *P < 0.05.

To determine whether the decreased expression of *CARMN* is responsible for SMC phenotypic modulation, we first transfected phosphorothioate modified antisense oligonucleotide (ASO) against *CARMN* or scrambled control into HCASMCs to knock down endogenous *CARMN* expression. EdU proliferation assays and Boyden migration assays revealed that knocking down *CARMN* significantly enhances cell proliferation and migration, respectively (**Figure 4A-C**). Moreover, depletion of endogenous *CARMN* significantly attenuated the expression of SM-contractile genes, including CNN1, ACTA2 and TGFB1I1, at both mRNA and protein levels (**Figure 4D-F**). These loss-of-function experiments suggest that decreased expression of *CARMN* plays a causal role in VSMC phenotypic switching from a contractile to a proliferative phenotype. Conversely, restoration of *CARMN* expression in HCASMCs mediated by *Carmn* adenoviral transduction promotes a contractile phenotype in VSMCs as indicated by increased numbers of spindle shaped HCASMCs (**Figure 4G & H**), decreased VSMC proliferation and migration (**Figure 4I-K**), and the up-regulation of SM-contractile genes at both mRNA and protein levels (**Figure 4L-N**). Of note, the *Carmn* adenovirus was constructed with transcript V2 that does not contain *Mir143/145* (**Figure 1I**), indicating that *Carmn* functions in SMCs is independent of the *Mir143/145* cluster. Collectively, these results suggest that *CARMN* is critical in maintaining a contractile phenotype in VSMCs.

**Figure 4.**
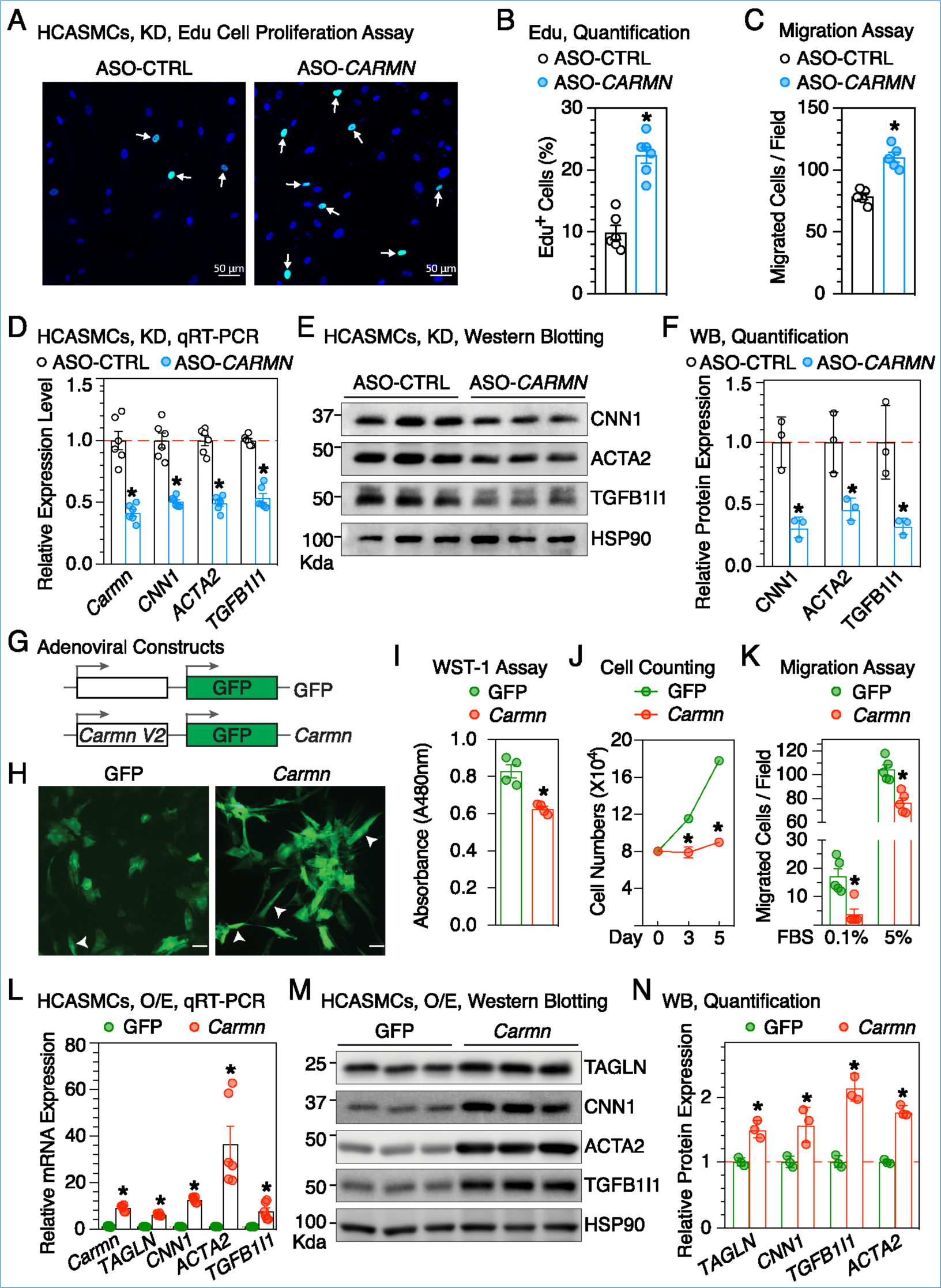
*CARMN* is required for maintaining the VSMC contractile phenotype. (**A**) *CARMN* or control phosphorothioate-modified antisense oligonucleotide (ASO) (ASO-CTRL) were transduced into human coronary artery SMCs (HCASMCs) for 5 days to knock down (KD) endogenous *CARMN* expression. EdU incorporation assays were performed to assess cell proliferation following depletion of *CARMN* in HCASMC. The arrows point to the representative EdU positive cells. (**B**) Quantitative analysis of EdU positive cells following *CARMN* depletion. N=5. *P <0.05. (**C**) Boyden chamber assays was performed to examine cell migration after knocking down *CARMN* in HCASMCs. N=6. *P <0.05. (**D**) Expression of SM-contractile genes was analyzed by qRT-PCR or (**E**) Western blot. (**F**) Densitometric analysis of protein levels as shown by Western blot (WB) in “**E**” normalized to the loading control HSP90 and relative to signals from ASO-CTRL group (set to 1). N=3. *P < 0.05. (**G**) *Carmn* or control GFP adenovirus were generated and transduced into HCASMCs for 48 hours. (**H**) An increased number of spindle shaped cells (arrowheads) was observed in *Carmn* over-expressed HCASMCs. Scale bar: 50μm. (**I**) Cell proliferation was measured by cell proliferation WST-1 kit and (**J**) cell number counting at the indicated time points. N=4 for WST-1 assay and N=3 for cell counting assay. *P < 0.05. (**K**) Over-expression of *Carmn* attenuated the migration of HCASMCs cultured in medium containing 0.1% and 5% FBS as revealed by Boyden chamber assays. N=5. *P < 0.05. (**L**) Expression of SM-contractile genes was analyzed by qRT-PCR or (M) Western blot. (**N**) Densitometric analysis of protein levels as shown in “**M**” normalized to the loading control HSP90 and relative to signals from GFP group (set to 1). N=3. *P < 0.05.

In conclusion, we have identified an evolutionarily conserved and SMC-specific lncRNA, *CARMN*, and confirmed the specificity *in vivo* using a GFP KI mouse model and RNA FISH assays. We also provided evidence showing *CARMN* is transcribed independent of *MIR143/145* and is expressed transiently in cardiomyocytes during early developmental stage and thereafter becomes restricted to SMCs in both human and mouse. We have demonstrated that *CARMN* is down-regulated in different vascular diseases and is a key regulator of VSMC phenotypic modulation. Since tissue/cell-specific genes are considered desirable for drug targets due to the reduced risk of side effects (32), *CARMN* may represent a promising therapeutic target for treatment of SMC-related diseases.

## Methods

Detailed methods are provided in the Supplemental Material.

*Study approval.* The use of experimental animals has been approved by the IACUC at Augusta University in accordance with NIH guidelines.

## Author contributions

JZ conceived and supervised the project. JZ and KD designed the experiments. KZ, JS, XH, LW performed the experiments. KB, RD and AV contributed to the RNA FISH assay. KZ analyzed data and wrote the manuscript. MX, ZZ, HX and JZ edited the manuscript.

## Acknowledgments

We thank Drs. Qing Lyu and Joseph Miano at Vascular Biology Center of Augusta University for helpful suggestions on RNA FISH assay in mouse tissues and sharing human *SENCR* Stellaris RNA probes. The work at the Zhou laboratory is supported by grants from the National Heart, Lung, and Blood Institute, NIH (R01HL132164 and R01HL149995). JZ is a recipient of Established Investigator Award (17EIA33460468) and Transformational Project Award (19TPA34910181) from the American Heart Association. KD is supported by postdoctoral fellowship from the American Heart Association (19POST34450071).

## Supplemental Methods

### Bioinformatics analysis of public RNA-seq or microarray datasets

Raw reads of RNA-seq data from 6 different human cell types, including vascular SMCs (VSMCs), endothelial progenitor cells, chondrocytes, human artery endothelial cells, human umbilical vein cells and lymphoblastoid cells (GSE100242) (1), mouse aortic tissues (GSE99318) (2), mouse brain, heart, liver, lung and testis (GSE61991) (3), were obtained from GEO database. Raw reads were first trimmed to remove low-quality reads using Trimmomatic 0.38 (4). By using TopHat version 2.1.0 (5), pass-filtered reads were then aligned to hg38 and mm10 genome for human and mouse RNA-seq data, respectively. Raw counts of each gene was estimated by featureCounts 1.6.2 (6) and Fragments Per Kilobase of exon per Million fragments mapped (FPKM) values (7) were calculated for these data sets. Transcripts Per Kilobase Million (TPM) values of whole genomic genes for 22 different human cell types were downloaded from ENCODE database (8) (**Supplemental Table I**). These data sets were used to identify human and mouse SMC/aorta-enriched lncRNAs by screening for lncRNAs with normalized expression value at least 1.5-fold higher in SMCs/aortic tissues than that in all the other types of cells/tissues.

To examine the distribution of *CARMN* expression in different types of vascular cells, count matrix of single cell RNA-seq (scRNA-seq) data for human coronary arteries from four cardiac transplant recipients was downloaded from GEO database (GSE131778) (9) and processed with R package “Seurat” for quality control, dimensionality reduction and clustering (10). Genes expressed in fewer than 5 cells, and cells expressing fewer than 500 or more than 3,500 genes, or containing >7.5% mitochondrial genes were removed. t-SNE plots were generated for visualization of different cell clusters. Normalized data was used to visualize the expression of selected genes in different cell types. The normalized expression table of scRNA-seq for normal mouse aorta was obtained from Broad Institute Single Cell Portal (https://singlecell.broadinstitute.org/single_cell/study/SCP289/single-cell-analysis-of-the-normal-mouse-aorta-reveals-functionally-distinct-endothelial-cell-populations) (11) and the t-SNE plots were generated using custom R script. Raw sequencing data and count matrix of scRNA-seq data of aortic root and descending aorta of mice at baseline before high-fat diet (HFD) as well as after 8 and 16 weeks of HFD, were obtained from GEO database (GSE131776) (9). STAR v2.7.1 (12) was used for demultiplexing the sequencing data and count matrix for cells from the three time points mentioned above were extracted and processed as described above in human scRNA-seq data analysis. To examine the distribution of *CARMN* expression in developmental and adult cardiac cells, count matrix of scRNA-seq data for cells isolated from embryonic mouse heart at E10.5 (GSE122403) (13), cardiomyocytes isolated from adult mouse heart by large-particle fluorescence-activated cell sorting (GSE133640) (14), and normal adult human heart (GSE109816) (15), were downloaded from GEO database. Count matrix of scRNA-seq for non-cardiac cells from adult mouse heart was obtained from ArrayExpress database(E-MTAB-6173) (16). Count matrix of scRNA-seq for developmental human heart at 6.5-7 post conception weeks (PCW) was downloaded from https://www.spatialresearch.org (17). The obtained count matrix or normalized expression matrix were processed with R package “Seurat” for quality control, dimensionality reduction and clustering (10).

BigWig files for ChIP-seq data of histone marks H3K27ac (ENCFF360FGJ and ENCFF656TFQ), H3K27me3 (ENCFF351CFT and ENCFF741TUI), H3K4me2 (ENCFF736JLA and ENCFF887SAU) and H3K4me3 (ENCFF302UFW and ENCFF948XBH) in human VSMCs and umbilical endothelial cells were obtained from ENCODE (8) and were used for visualization of human *CARMN* transcription activity with Integrative Genomics Viewer (IGV) (18).

Microarray data of intact tissues and atheroma plaque of carotid endarterectomy from 32 hypertensive patients was analyzed using NCBI online GEO2R tool (GSE43292) (19). Raw RNA-seq data of human cerebral aneurysms as well as healthy superficial temporal arteries (GSE66240) (20), human saphenous vein VSMCs in the absence or presence of platelet-derived group factor (PDGF-BB) or Interleukin-1*α* stimulation (GSE69637) (21), and rat VSMCs with or without PDGF-BB treatment for 24 hours (GSE111714), were downloaded from GEO database. The rat RNA-seq data was first used to de novo assemble the rat VSMC transcriptome with Cufflinks (22) and the assembled transcripts that include *Carmn* gene were used for gene expression analysis. The other RNA-seq data were re-analyzed using the RNA-seq data analysis method described above.

### Quantitative reverse transcription-PCR (qRT-PCR) analysis

Total RNA from tissues or cells was isolated with TRIzol reagent (Invitrogen). 0.8μg of RNA was used as template for reverse transcription (RT) with random hexamer primers using the High Capacity RNA-to-cDNA Kit (Invitrogen). Real time PCR was performed with respective primers listed in **Supplemental Table II**. All samples were amplified in duplicates. Relative gene expression was converted using the 2^-êêCT^ method against the internal control house-keeping gene hypoxanthine phosphoribosyltransferase 1 (*Hprt*) or *Rplp0* (for rat) where êêCT= (CT_experimental gene_ - CT_experimental Hprt_) - (CT_control gene_ - CT_control Hprt_). The relative gene expression in control group was set to 1.

### Protein extraction and Western blotting

Protein lysates were extracted from tissues or cells by RIPA buffer (Thermo Fisher Scientific) plus 1% protease/phosphatase inhibitor cocktail (Thermo Fisher Scientific). After sonication and centrifugation of the tissue or cell lysates, proteins in the supernatant were quantified by BCA assay (Thermo Fisher Scientific) and resolved on a 7.5%,10% or 12.5% SDS-PAGE gel at 10μg per lane as appropriate. Primary antibodies used in this study were: CNN1 (Sigma, C2687, mouse 1:1000), TAGLN (Abcam, ab10135, goat, 1:2000), TGFB1I1 (BD, 611164, mouse, 1:5000), ACTA2 (Sigma, A2547, mouse, 1:5000), HSP90 (Cell signaling, 4874, rabbit, 1:1000) and GFP (Abcam, ab290, rabbit, 1:2500). Secondary antibodies conjugated with horseradish peroxidase were then used to visualize the target proteins in blot. Image was acquired by ImageQuant LAS 4000 Imaging Station (GE) and band densities were quantified using the ImageJ software.

### Rapid amplification of cDNA ends (RACE)

Total RNA of mouse thoracic aorta was used for RACE to define 5’ and 3’ ends of mouse *Carmn* transcripts using 5’/3’ RACE kit (Sigma-Aldrich) according to the manufacturer’s instructions. The obtained full-length *Carmn* V2 and V3 transcripts were cloned into pSC-B blunt vector (Stratagene) for Sanger sequencing. The full-length sequence of *Carmn* V2 and V3 obtained by RACE have been deposited into GenBank under the accession number of MN904529 and MN904530, respectively.

### Identification of *Carmn* homolog in rat

The full-length sequence of mouse *Carmn* V2 transcript was used as query sequence for BLAST against rat genome to determine the *Carmn* homologous region in rat genome. Forward and reverse primers specifically targeting rat sequence within the homologous region spanning different exons (**Online Table II** and **Supplemental Figure 1E**) were designed and used for PCR amplification with rat aorta cDNA as template. The PCR product was subsequently cloned into pSC-B blunt vector (Stratagene) for Sanger sequencing and the obtained sequence were mapped back to rat genome for confirmation.

### Generation of *Carmn* KI reporter mice

The *Carmn* KI mouse line with a promoterless, inverted splicing acceptor (SA)-membrane bound GFP cassette flanked by two pairs of *LoxP* sites and *Lox2272* sites in an opposite orientation was generated by Cyagen (www.cyagen.com) using the homologous recombination strategy. The cassette was placed in the intron between exon 2 and exon 3 of *Carmn* gene locus (Figure 2A). Briefly, mouse genomic fragments containing 5’ and 3’ homology arms of *Carmn* gene were amplified from C57BL/6 BAC clone with high fidelity Taq DNA polymerase and were sequentially assembled into a targeting vector containing *LoxP* and *Lox2272* recombination sites, *neomyosin* (neo) cassette flanked by a pair of *Rox* sites, inversed SA-membrane bound GFP-PolyA cassette and a DTA negative selection marker. The final targeting vector was sequenced to ensure the DNA integrity and linearized with NotI prior to electroporation into C57BL/6 ES cells. After selected with G418, DNA was extracted from the positive ES cell clones and subjected for long range PCR followed by Southern blot to confirm the correct homologous recombination. The *Carmn* targeted ES cells were then injected into C57BL/6 host blastocysts to generate chimera mice. The resulted chimera mice were bred with wild-type (WT) C57BL/6J mice to obtain germline transmission WT/neo mice. WT/neo female mice were further crossed to male Dre deleter mice to remove the neo cassette. The resulting mice were intercrossed and maintained as *Carmn* conditional PFG mice on C57BL/6J background. Cre-mediated recombination will first cause inversion of the reversed GFP cassette (PFG to GFP), resulting in GFP being in the correct 5’-3’ orientation relative to the endogenous *Carmn* transcript. This also places the pair of *LoxP* sites or the pair of *Lox2272* at the same orientation thus allowing Cre to further excise the genomic region including *Carmn* exon 2. As a result, *Carmn* transcription will result in splicing of *Carmn* exon 1 to the GFP gene trap cassette, leading to GFP expression driven by the endogenous *Carmn* gene promoter (referred to as GFP, G mice). GFP expression will therefore faithfully display the endogenous *Carmn* expression *in vivo* (Figure 2A). To visualize the *Carmn* expression *in vivo*, *Carmn*^GFP/WT^ (GW) mice were generated by crossing female *Carmn*^PFG/WT^ (PW) with male mice carrying *CMV*-*Cre* (JAX #: 006054) which ubiquitously express Cre in all mouse tissues.

### Sections, directly visualizing GFP and immunofluorescence (IF)

Aortic tissues from adult mice or embryos (E16.5) of GW genotype were harvested and fixed with 4% paraformaldehyde on ice for 30 minutes. After washing with ice-code phosphate-buffered saline (PBS) twice for 10 minutes each time, tissues were then soaked in 30% sucrose/PBS on a shaker in 4°C overnight until the tissues settled down. The processed tissues were subsequently embedded in optimal cutting temperature medium (OCT) and cryosections were prepared at 8-μm thickness. For directly visualizing GFP, OCT sections were air dried for 10-15 minutes then washed with PBS (3 times x 5 min/wash) to remove OCT. The slides were then briefly air dried and covered with Prolong Gold DAPI mounting medium (Invitrogen) to counter-stain nuclei. GFP and DAPI signals were then acquired by a confocal microscopy (LSM 780 upright, Zeiss) using 488, 405-nm excitation channels, respectively. For IF staining of GFP, the SM-enriched marker ACTA2, and the endothelial marker PECAM1 in aortic tissues, antigen retrieval was carried out by microwaving to heat at 98°C for 10 minutes in citric acid buffer (10 mM, pH 6.0) to quench all of endogenous fluorescent signal. After blocking with goat serum (10%, Invitrogen) for 30 minutes, sections were then incubated with anti-ACTA2 (Sigma, A2547, mouse, 1:200), anti-GFP (Abcam, ab290, rabbit, 1:200), and anti-PECAM1 (BD Pharmingen, 550274, rat, 1:30) antibodies. Subsequently, sections were incubated with secondary antibodies against mouse, rabbit or rat that is conjugated with different fluorophore for 1 hour at room temperature. Following 3 washes with PBS, sections were then immersed with mounting medium (ProLong Gold anti-fade reagent with DAPI, Invitrogen) to stain nuclei. Images were collected by using a confocal microscopy (LSM 780 upright, Zeiss) at the imaging core of Augusta University.

### RNA fluorescence *in situ* hybridization (FISH) in mouse aorta

Immediately after dissection, aorta from adult WT mouse was embedded in OCT and 8-μm thick cryosections were prepared. Stellaris RNA FISH probes for *Carmn* were designed and purchased from Biosearch Technologies. RNA FISH were performed following a protocol adapted from the manufacture’s instruction for frozen tissues. Briefly, OCT sections were air dried for 10-15 min and then fixed with 3.7% formaldehyde (Sigma). After fixation, sections were washed twice with 1x PBS and permeabilized with 0.2% Triton X-100. Slides were then immersed in 70% ethanol for 3 hours at room temperature, followed by rinses in 90% and 100% ethanol for 3 min for dehydration. After hybridization at 37°C overnight in Stellaris RNA-FISH hybridization buffer (#SMF-HB1-10, Biosearch) with *Carmn* probes, sections were washed in 2x saline-sodium citrate (SSC) and 10% formamide for 30 minutes at 37°C and then immersed with mounting medium (ProLong Gold anti-fade reagent with DAPI, Invitrogen) to stain nuclei. Hybridization with only hybridization buffer or human specific *SENCR* Stellaris probe (23) was used as negative controls.

### RNA FISH in mouse VSMCs

For RNA FISH in mouse VSMCs, the full-length mouse *Carmn* transcript V2 was obtained by PCR using mouse aortic cDNA and primers listed in **Supplemental Table II**. The PCR product was then ligated into pSC-B blunt vector (Stratagene) and the orientation of the inserts was determined by Sanger sequencing. Digoxigenin labelled sense or antisense RNA probes were then synthesized by MAXIscript T7 in vitro transcription kit (Ambion, Austin, TX) using T7 promoter. FISH assay was performed essentially following the protocol previously reported (24). Due to the undetectable level of *Carmn* expression in mouse VSMCs cultured in complete medium, cells transduced with *Carmn* adenovirus were used for RNA FISH. Briefly, mouse VSMCs were cultured in 4-well chamber slides and transduced with *Carmn* adenovirus. After adenoviral transduction for 48 hours, cells were fixed in 4% paraformaldehyde, permeabilized in a 1:1 solution of acetone and methanol, and then hybridized with digoxigenin-labelled *Carmn* sense or antisense RNA probes. After peroxidase quenching and blocking, the hybridized cells were incubated with peroxidase-conjugated anti-digoxigenin antibody (Roche, Indianapolis, IN, USA) and revealed using SuperGlo^TM^ Green Immunofluorescence Amplification Kits (Fluorescent Solutions, Augusta, GA, USA). Nuclei were then counter-stained with DAPI (Invitrogen, Carlsbad, CA, USA). RNA-FISH images were collected by using a confocal microscopy (LSM 780 upright, Zeiss) at the imaging core of Augusta University.

### Mouse carotid artery ligation and femoral artery wire injury

Mouse carotid ligation and femoral wire injury was performed as we previously described (25, 26). Briefly, mice were anesthetized by intraperitoneal injection of a mixed solution of xylazine (5 mg/kg body weight) and ketamine (80 mg/kg body weight). For carotid ligation injury, the left common carotid artery was dissected and completely ligated just proximal to the carotid bifurcation. The right carotid artery served as an uninjured contralateral control. The femoral artery wire injury was induced by inserting a spring wire (0.38 mm in diameter; Cook Medical) from an exposed muscular branch artery into the left femoral artery (LFA) for > 5 mm toward the iliac artery. The wire was left in place for 1 minute to denude and dilate the femoral artery. Then, the wire was removed, and the proximal portion of the muscular branch artery was secured with silk sutures followed by restoration of blood flow in the LFA. The right femoral artery (RFA) was not subjected to wire injury and therefore served as a contralateral control.

### Rat carotid artery balloon injury

Rat carotid artery balloon injury was performed as we previously described (24-26). Briefly, male Sprague-Dawley rats (350 g; Taconic Farms, Germantown, NY) were anesthetized with ketamine (70 mg/kg) and xylazine (4.6 mg/kg) via intraperitoneal injection. Following a midline cervical incision and muscular tissues separation, the left common carotid artery was exposed and blunt dissection was performed alongside the artery by dull forceps to expose the carotid artery bifurcation into the internal/external branches. Blood flow cessation was achieved by arterial clamps and a small arteriotomy was made in the external carotid artery. A 2F Fogarty balloon embolectomy (Edwards) was inserted through the small cut and passed into the common carotid artery. After balloon inflation at 1.6-2.0 atmospheric pressure, the catheter was partially withdrawn and reinserted 3 times. After the catheter was removed, the external carotid artery was secured with silk sutures, and the blood flow in the common carotid artery and its internal branch was restored by releasing arterial clamps. The intact right carotid artery served as contralateral control.

### Rat VSMC culture and PDGF-BB treatment

Rat aortic tissues were harvested and rat primary aortic SMCs were prepared and cultured as described in our previous studies (26-29). Briefly, male Sprague-Dawley rats (200-250 grams, Taconic Farms, Germantown, NY) were euthanized by CO2 and the thoracic aorta was dissected to remove adhering periadventitial tissue and the endothelium was denuded with a catheter. The aorta was digested with a Blendzyme III solution (Roche, 0.5U/ml) for 10 min at 37°C followed by dissection to remove the adventitial layer, then the remaining medial layer was minced into small pieces for a second digestion with Blendzyme III for 2 hours at 37°C. Following removal of digestion solution and re-suspending in 10% FBS DMEM medium, cells were gently liberated with a pipette and transferred into culture dishes. Every batch of smooth muscle cells was tested by smooth muscle maker ACTA2 staining to ensure the purity of primary VSMCs above 95% and used for experiments before passage 6. For PDGF-BB treatment experiments, rat primary aortic VSMCs were grown to 80-90% confluence and serum-starved for 24 hours and then treated with recombinant rat 50 ng/ml PDGF-BB (platelet-derived growth factor BB, Calbiochem) for 24 hours as described in our previous reports (26, 28). Cells treated with vehicle served as control. Following PDGF-BB or vehicle administration cells were harvested for total RNA to detect rat *Carmn* expression by qRT-PCR.

### Cell culture for HCASMCs (human coronary artery smooth muscle cells)

HCASMCs (Gibco, Cat. #: C-017-5C; Lot #: 1130140 and Lot #: 1689414), vascular cell basal medium (Cat. #: M231500) and smooth muscle growth supplement with 5% FBS (SMGS, Cat. #: S00725) were purchased from Thermo Fisher Scientific. HCASMCs (passages 3-5) were cultured incubated in vascular cell basal medium containing SMGS (complete medium) and 5.5 mM D-glucose along with antibiotic/antimycotic solution in a humidified atmosphere of 95% air and 5% CO_2_ at 37°C as we recently reported (30). Sub-confluent HCASMCs were trypsinized, centrifuged, and seeded onto petri-dishes or multi-well plates. In some experiments, HCASMCs differentiated status was induced by substituting the complete medium with vascular cell basal medium without SMGS for 48 hours before treatment. To compare *CARMN* expression between VSMCs under differentiation and de-differentiation status, HCASMCs were cultured in SMC differentiation (Cell Applications, Cat. #: 311D-250) medium and growth medium (Cell Applications, Cat. #: 311K-500) for 48 hours, and harvested for qRT-PCR analysis.

### Antisense oligonucleotides (ASO) transfection

Three phosphorothioate modified ASOs targeting the common region of human *CARMN* transcripts (*CARMN*-ASO-1, 5’-TCTGTGAAAGGTGATG-3’; *CARMN*-ASO-2, 5’-CAGAGTTCTTGCTTCTCTGA-3’; *CARMN*-ASO-3, 5’-AGGTTCCACTTCTTAACGAG-3’) and a scrambled ASO (5’-TCATACTATATGACAG-3’) which was used as control were synthesized by Exiqon. Delivery of ASO (mixed *CARMN*-ASO-1, 2 and 3) into primary human HCASMCs were carried out by using Lipofectamine RNAiMAX Transfection Reagent (Invitrogen, Cat. #: 13778) at a final concentration of 25nM. Cells were harvested for EdU assay, Boyden chamber migration assay, qRT-PCR and Western blot analysis after 5 days post transfection.

### 5-Ethynyl-2-deoxyuridine (EdU) incorporation assay

Edu incorporation assay was performed using the Click-iT EdU imaging kit (Thermo Fisher Scientific, Cat. #: C10339) according to the manufacturer’s instructions. Briefly, following knocking down of *CARMN*, HCASMCs were incubated with EdU (10 mM) in vascular cell full medium overnight. EdU-positive nuclei were detected following the manufacturer’s protocol and imaged using a confocal microscopy (780 upright, Zeiss).

### Boyden chamber migration assay

Boyden chamber assays were performed as previously described (24, 25, 29). Briefly, following knock-down or over-expression of *CARMN*, HCASMCs were grown in basal medium for 48 hours. Subsequently, the cells were trypsinized and seeded into Boyden chambers (PET track-etched, 8-um pores, 24-well format; Becton Dickinson) in basal medium. Chambers were then immersed in basal or full medium (5% FBS) for 5 hours. The top side of the membranes was swabbed to remove cells, and then cells on the bottom surface of the membrane were fixed with 4% paraformaldehyde, stained with DAPI to visualize nuclei, and counted under a fluorescence microscopy. Five identically located fields per membrane were averaged for quantification of migrated cell numbers.

### Adenoviral construction and cell infection

The full-length mouse *Carmn* transcript V2 determined by RACE was amplified by PCR and ligated into AdTrack shuttle vector. As this vector contains an independent cytomegalovirus (CMV) promoter-driven transcription cassette for green fluorescent protein (GFP), the efficiency of transduction was directly monitored by visualization of GFP expression. Transferring the *Carmn* and GFP expression cassettes into AdEasy viral backbone, viral packaging, and cell transduction was performed as we previously described (30). Adenovirus expressing *Carmn* or GFP was added into HCASMCs cultured in full medium for 48 hours and cells were used for WST-1 proliferation assay, cell counting, Boyden chamber migration assay, qRT-PCR and western blot analysis. The adenovirus expressing GFP alone served as control.

### WST-1 proliferation assay

The assay was performed according to manufacturer’s instructions as we previously reported (26). Briefly, following over-expression of *Carmn*, HCASMCs were maintained in the basal medium for additional 48 hours to allow growth arrest. After 24 hours, WST-1 reagent (Sigma-Aldrich, Cat. #: 5015944001, 10μl / 100μl of media) was added to the culture medium and cells were kept at 37°C for 2 hours. Afterwards, the absorbance at 450 nm, as a direct readout of cell proliferation, was measured using a microplate reader.

### Cell counts

Sub-confluent HCASMCs were equally plated onto 6-well plates and were transduced with GFP or *Carmn* adenovirus. During the treatment period, the medium was replaced every 48 hours with a fresh complete medium. After 3 or 5 days of infection, VSMCs were trypsinized and the changes in cell numbers were determined using a hemocytometer

### Statistical analysis

All data are expressed as means ± SE of at least three independent experiments. Statistical analysis of the data was performed using one-way analysis of variance (ANOVA) followed by Bonferroni *t*-test for data involving more than two groups. Unpaired two tailed *t*-test or paired two tailed *t*-test was used for data involving two groups only (GraphPad). Values of p *≤* 0.05 were considered statistically significant.

---

**Supplemental Figure 1.**
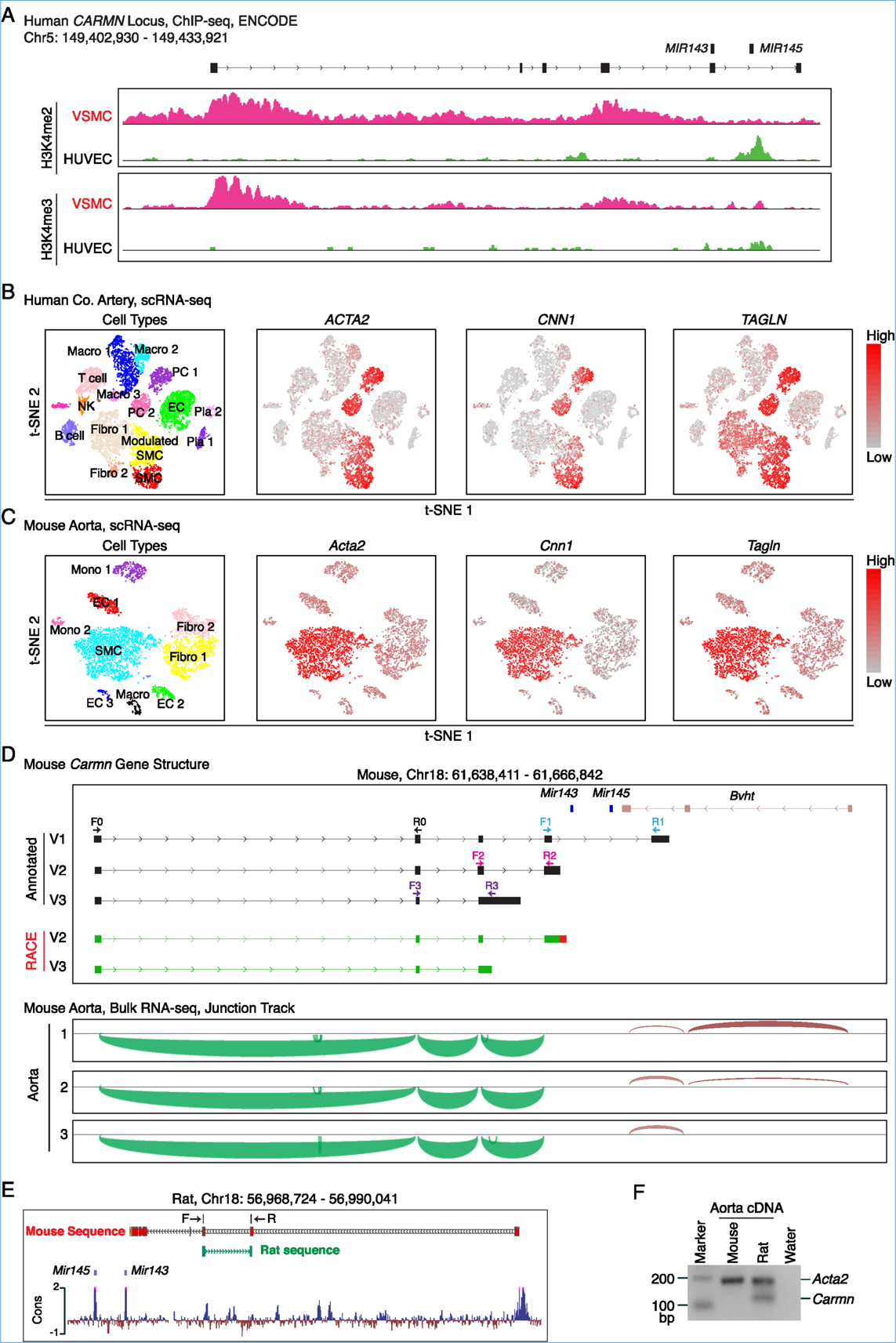
D**e**termination **of the expression and structure of human, mouse and rat *CARMN* gene. (A)** ChIP-seq tracks of H3K4me2 and H3K4me3 in human *CARMN* locus in human VSMC vs HUVEC as revealed by the ENCODE project. **(B)** t-SNE visualization of cell types isolated from the human right coronary artery of four cardiac transplant recipients (left), and the expression of *ACTA2*, *CNN1*, and *TAGLN* revealed by scRNA-seq. The color scale on the right indicates the expression levels of these genes. Fibro: fibroblast; EC: endothelial cell; PC: pericyte cell; Macro: macrophage; NK: natural killer cell; Pla: plasma cell. **(C)** t-SNE visualization of cell types present in normal mouse aorta (left), and the expression of *Acta2, Cnn1, Tagln* revealed by scRNA-seq. The color scale on the right indicates the expression levels of these genes. Mono: monocyte. **(D)** RNA-seq of 3 mouse aortic tissues reveals the annotated *Carmn* V1 transcript was undetectable in mouse aorta. The upper panel shows the annotated and RACE-identified mouse *Carmn* gene structure. The bottom Sashimi plots show the alternative splicing junctions of *Carmn* gene. F0/R0: primers used for qPCR analysis of all *Carmn* transcripts. F1/R1: primers used for qRT-PCR analysis of *Carmn* transcript V1. F2/R2: primers used for qRT-PCR analysis of *Carmn* transcript V2. F3/R3: primers used for qPCR analysis of *Carmn* transcript V3. The sequence of these primers was listed in Supplemental Table II. **(E)** Determination of rat *Carmn* homolog. The mouse *Carmn* transcript V2 sequence was used as a query sequence to perform BLAST against rat genome and a putative rat *Carmn* homolog positionally conserved to *miR143/145* locus, was obtained (Mouse sequence, red). A pair of primers (F: forward; R: reverse) specifically targeting rat genome and spanning two different putative exons within the homologous region of rat *Carmn* gene were designed and used for PCR amplification with rat aorta cDNA as template. The obtained sequence was subjected to Sanger sequencing and mapped back to rat genome by BLAST for confirmation (Rat sequence, green). (**F**) Gel picture of PCR product showing that PCR with rat specific primers indicated in “**E”** (F and R) specifically generated a product from rat aorta cDNA, but not from mouse aorta cDNA or water. Primers for *Acta2* of both rat and mouse was used as an internal control for PCR and water was used as a negative control.

**Supplemental Figure 2.**
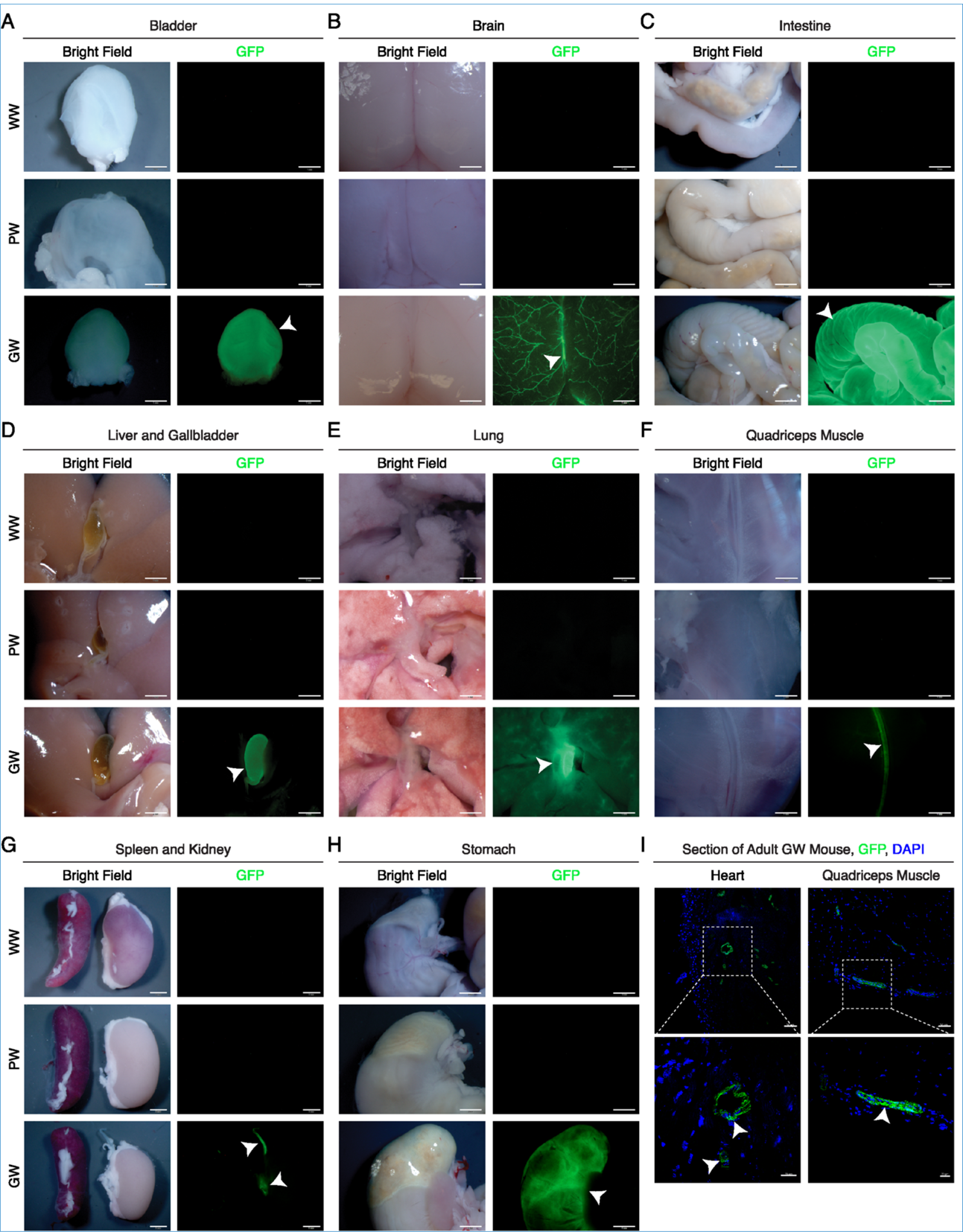
D**i**rect **visualization of GFP expression in a variety of tissues from mice with different genotypes.** Different tissues including **(A)** bladder, **(B)** brain, **(C)** intestine, **(D)** liver and gallbladder, **(E)** lung, **(F)** quadriceps muscle, **(G)** spleen and kidney, and **(H)** stomach, were dissected from WW, PW and GW mice and photographed under bright field or epifluorescence (GFP channel). Arrowheads point to GFP positive tissues. Arrowhead in “**D**” points to gallbladder. Arrowhead in “**E**” points to bronchus/pulmonary artery. The upper and bottom arrowheads in “**G**” point to the ureter and renal artery, respectively. **(I)** Visualization of GFP expression in sections of heart (left) and quadriceps muscle (right) showing GFP is specifically expressed in SMCs within the blood vessels of these tissues, but not in cardiomyocytes or skeletal muscle cells. Cell nuclei were stained with DAPI (blue). Arrowheads point to GFP signal. Scale bar: 1 mm in “**A**”, “**B**”, “**E**”, and “**F**”; 2 mm in “**C**”, “**D**”, “**G**”, and “**H**”; 50 and 20 μm in upper and bottom panels in “**I**”, respectively.

**Supplemental Figure 3.**
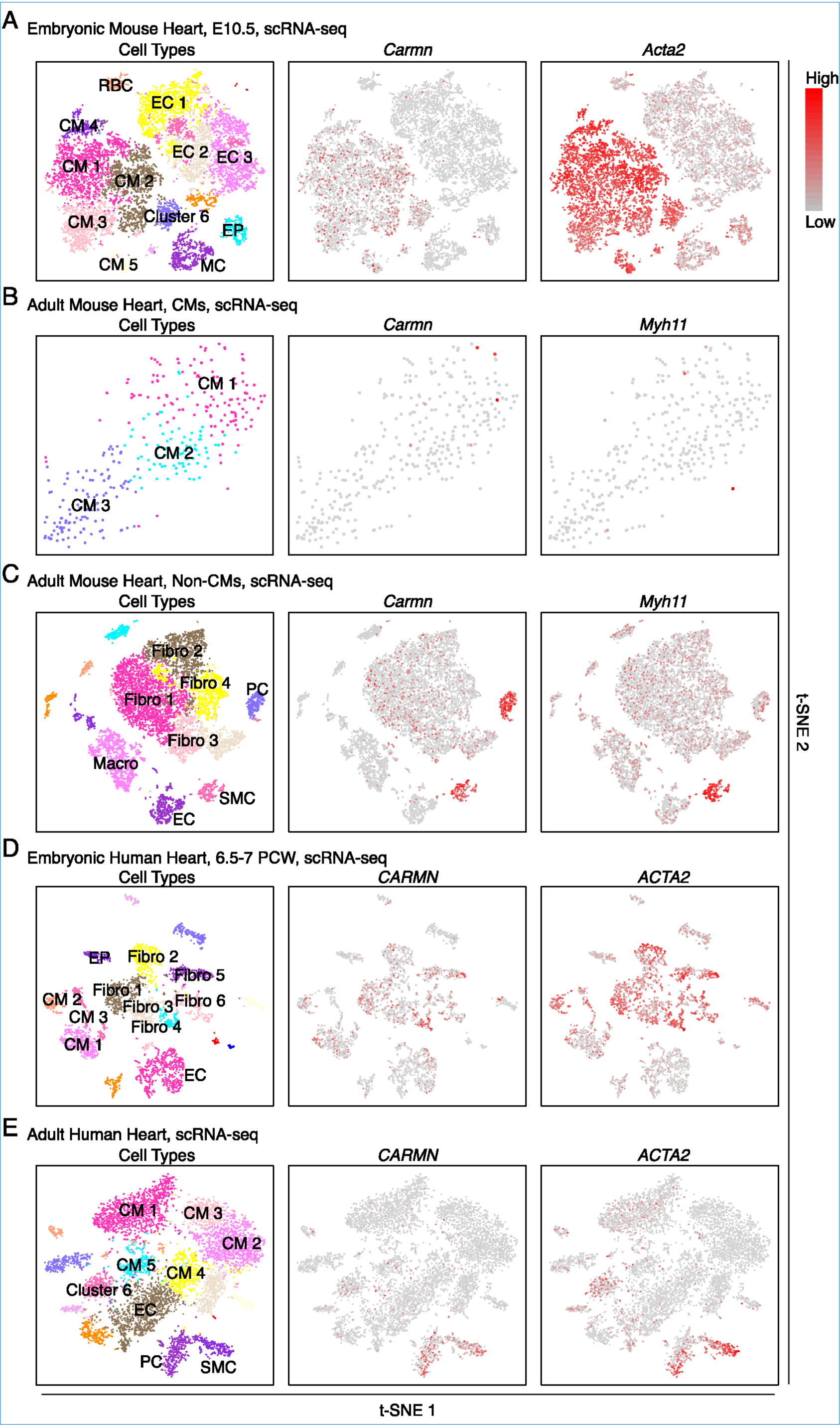
scRNA**-seq analysis of embryonic and adult hearts in both mouse and human.** t-SNE visualization of cell types and expression of *Carmn* and the SM marker *Acta2* or *Myh11* as indicated in **(A)** embryonic E10.5 mouse heart, **(B)** cardiomyocytes (CMs) isolated from adult mouse heart, **(C)** non-cardiomyocytes isolated from adult mouse heart, **(D)** embryonic 6.5-7 post conception weeks (PCW) human heart, and **(E)** adult human heart, as revealed by scRNA-seq. In E10.5 mouse heart **(A)**, *Carmn* is slightly expressed in a novel cell population expressing both cardiomyocyte and endothelial markers (Cluster 6). In adult human heart **(E)**, a novel subset of cells expressing both cardiomyocyte and SMC markers such as *ACTA2* (Cluster 6) display some extent expression of *CARMN*. EC: endothelial cell; EP: epicardial cell; MC: mesenchymal cell; RBC: red blood cell; Fibro: fibroblast; SMC: smooth muscle cell; PC: pericyte; Macro: macrophage. The color scale on the right indicates the expression levels of these genes.

**Supplemental Figure 4.**
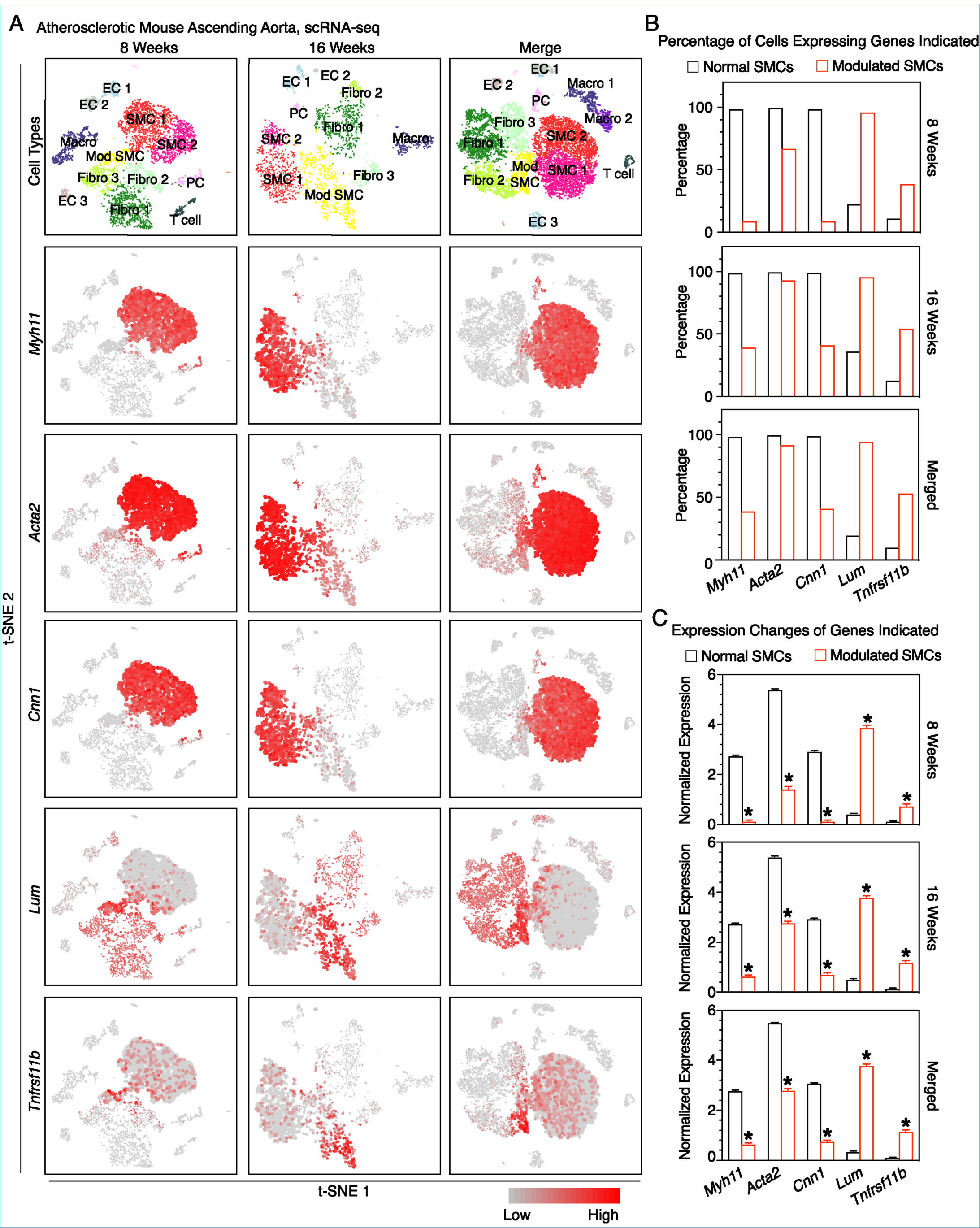
E**x**pression **of SMC differentiation and de-differentiation markers in normal and modulated SMCs in atherosclerotic mouse aorta revealed by scRNA-seq. (A)** t-SNE visualization of cell types present in the mouse aortic root and descending aorta after 8 (upper, left) and 16 weeks (upper, middle) of HFD, and merged cells from all time points including baseline before HFD feeding, after 8 and 16 weeks of HFD feeding (upper, right), and the expression of SMC differentiation markers including *Myh11*, *Acta2*, *Cnn1*, and de-differentiation markers including *Lum* and *Tnrfsf11b*. SMC: smooth muscle cell; Fibro: fibroblast; EC: endothelial cell; PC: pericyte; Macro: macrophage. The color scale on the bottom indicates the expression levels of these genes. (B) Quantitative analysis of percentage and **(C)** expression level of SMC differentiation (*Myh11, Acta2, Cnn1*) and de-differentiation (*Lum, Tnfrsf11b*) markers in normal and modulated SMC clusters revealed in “**A**”. *FDR <0.05.

**Supplemental Table I.** Information of human cell types from ENCODE used for identifying VSMC-specific lncRNAs

| *Cell type* | *ENCODE ID* | *GEO ID* | *Sample ID* |
| --- | --- | --- | --- |
| Coronary artery SMCs | ENCSR000AAG | GSE78534 | GSM2072316 |
|  |  |  | GSM2072317 |
| Aortic artery SMCs | ENCSR000AAA | GSE78528 | GSM2072304 |
|  |  |  | GSM2072305 |
| Umbilical artery SMCs | ENCSR000AAU | GSE78548 | GSM2072344 |
|  |  |  | GSM2072345 |
| Pulmonary artery SMCs | ENCSR000AAN | GSE78541 | GSM2072330 |
|  |  |  | GSM2072331 |
| Hepatocytes | ENCSR859HWB | GSE88601 | GSM2344260 |
|  |  |  | GSM2344261 |
| Cardiac fibroblasts | ENCSR369RVN | GSE78645 | GSM2072536 |
|  |  |  | GSM2072535 |
| Bronchus fibroblast | ENCSR620NSN | GSE78660 | GSM2072565 |
|  |  |  | GSM2072566 |
| Glomerular endothelial cell | ENCSR878EUT | GSE88615 | GSM2344284 |
|  |  |  | GSM2344285 |
| Microvascular endothelial cells | ENCSR815UVL | GSE78671 | GSM2072587 |
|  |  |  | GSM2072588 |
| Vein endothelial cells | ENCSR000CUG | GSE78609 | GSM2072464 |
|  |  |  | GSM2072465 |
| Thoracic aortic endothelial cells | ENCSR000CUK | GSE78613 | GSM2072472 |
|  |  |  | GSM2072473 |
| Umbilical vein endothelial cells | ENCSR000COZ | GSE78588 | GSM2072423 |
|  |  |  | GSM2072424 |
| Pulmonary artery endothelial cells | ENCSR000AAM | GSE78540 | GSM2072328 |
|  |  |  | GSM2072329 |
| Coronary artery endothelial cells | ENCSR000AAF | GSE78533 | GSM2072314 |
|  |  |  | GSM2072315 |
| Blood vessel endothelial cells | ENCSR000AAI | GSE78536 | GSM2072320 |
|  |  |  | GSM2072321 |
| Cardiomyocytes | ENCSR294NDO | GSE88167 | GSM2343523 |
|  |  |  | GSM2343524 |
| B cells | ENCSR000CTV | GSE90268 | GSM2400232 |
|  |  |  | GSM2400233 |
| Monocytes | ENCSR000CUC | GSE90272 | GSM2400240 |
|  |  |  | GSM2400241 |
| Skin melanocyte 1 | ENCSR000CUR | GSE78619 | GSM2072483 |
|  |  |  | GSM2072484 |
| Skin melanocyte 2 | ENCSR000CUQ | GSE78618 | GSM2072481 |
|  |  |  | GSM2072482 |
| Skeletal muscle myoblast 1 | ENCSR444WHQ | GSE78649 | GSM2072543 |
|  |  |  | GSM2072544 |
| Skeletal muscle myoblast 2 | ENCSR000COY | GSE78587 | GSM2072421 |

**Supplemental Table II.**
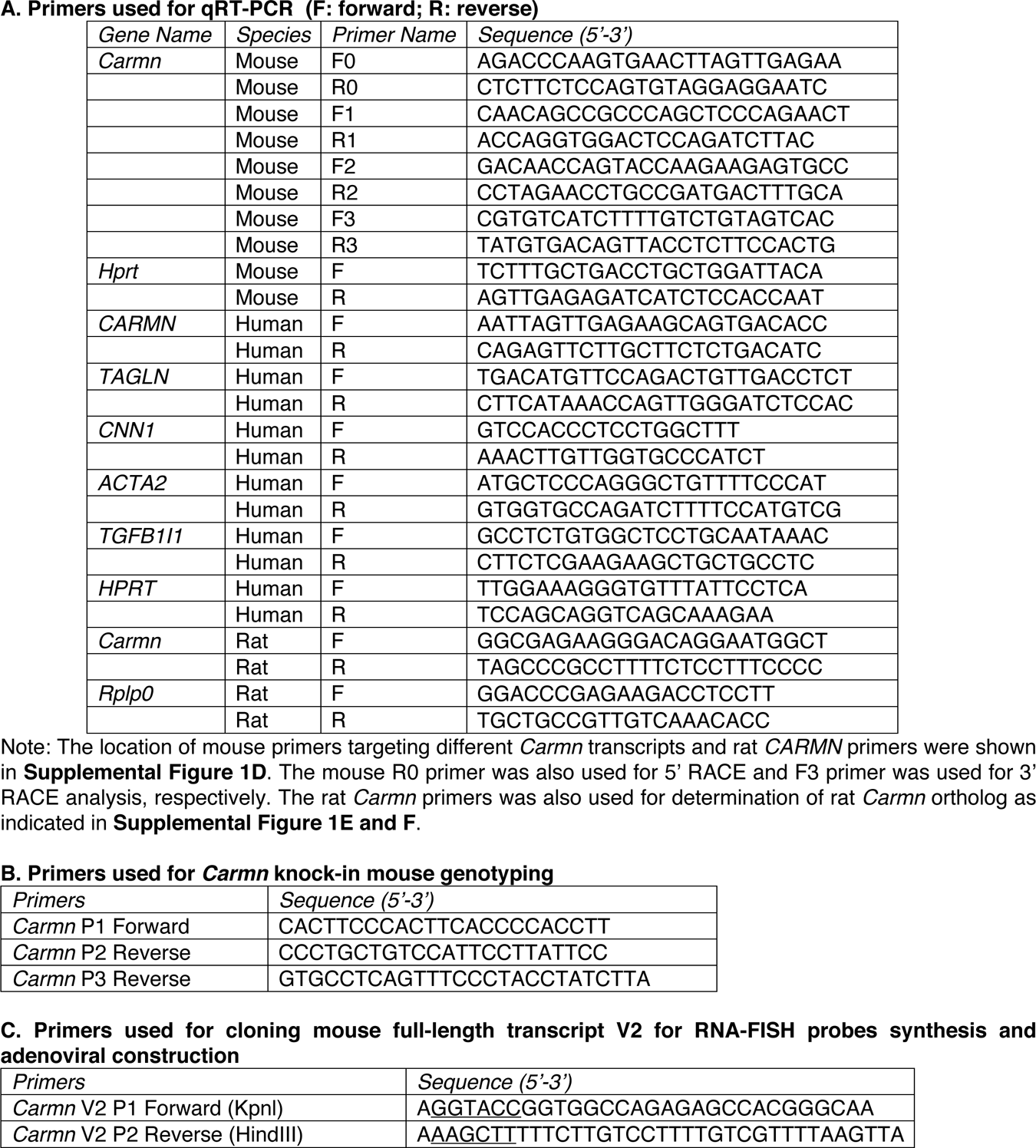
List of oligonucleotides used in this study

**Original Western blot pictures for Figure 2D.**
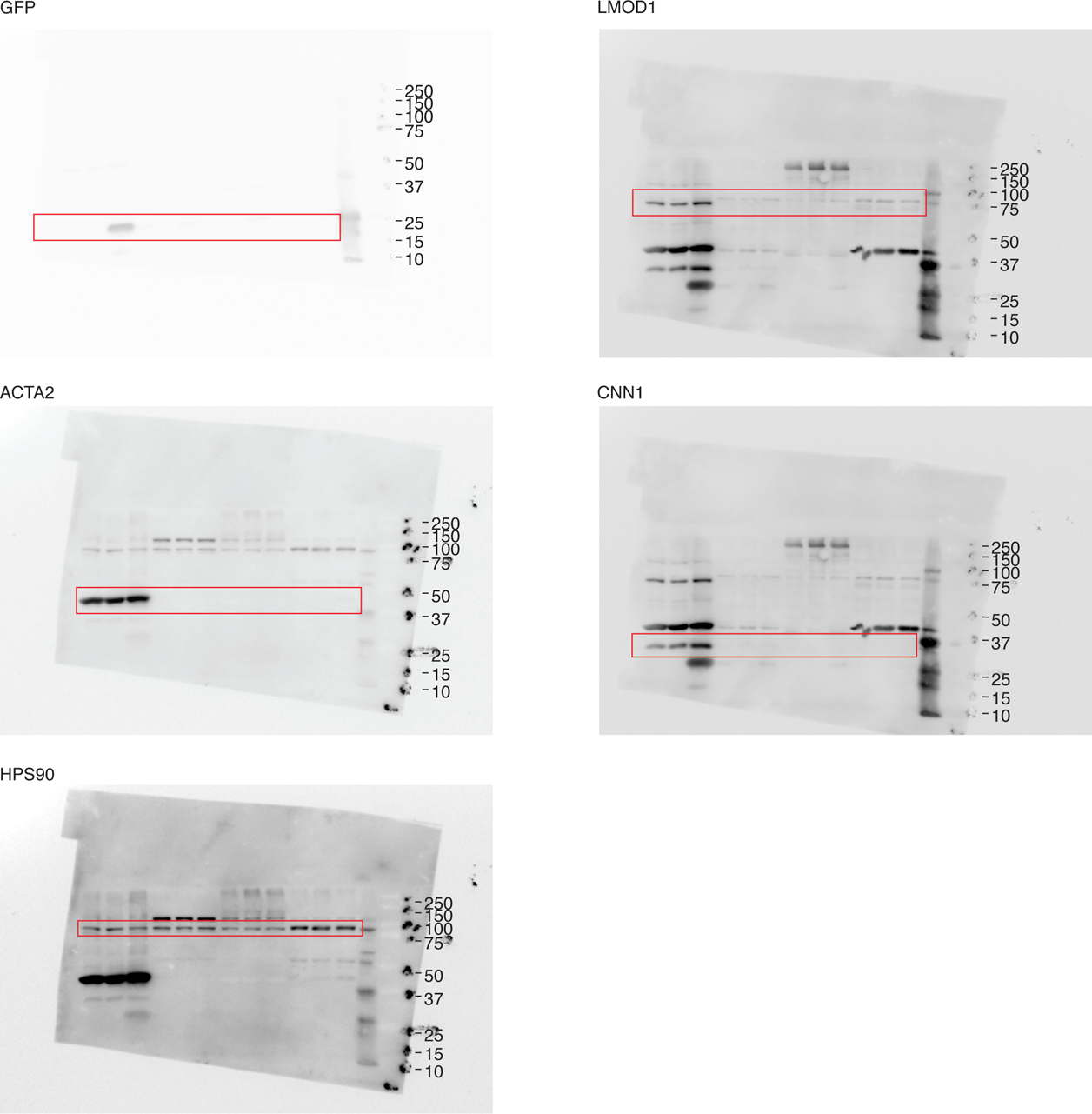

**Original Western blot pictures for Figure 4E.**
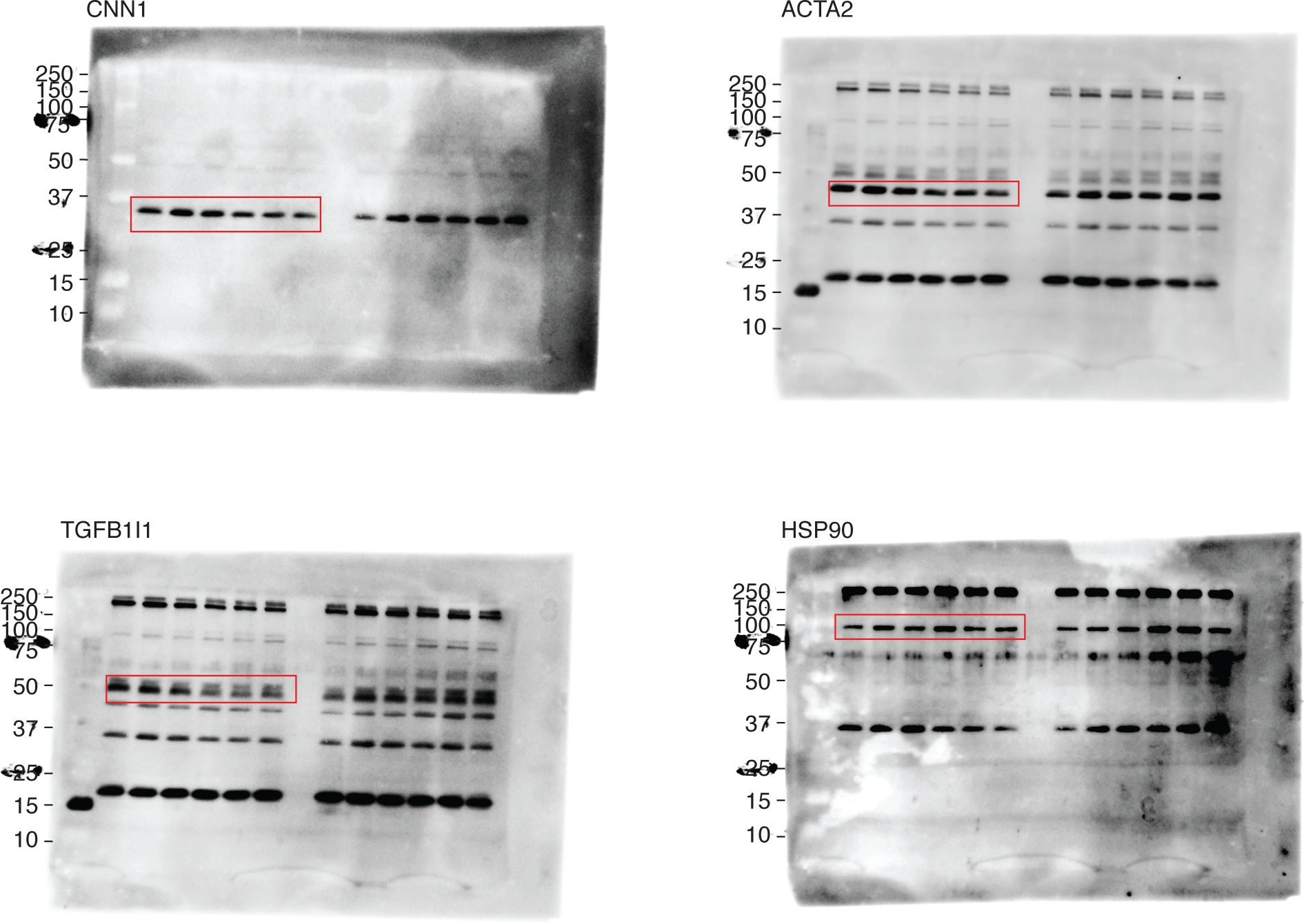

**Original Western blot pictures for Figure 4M.**
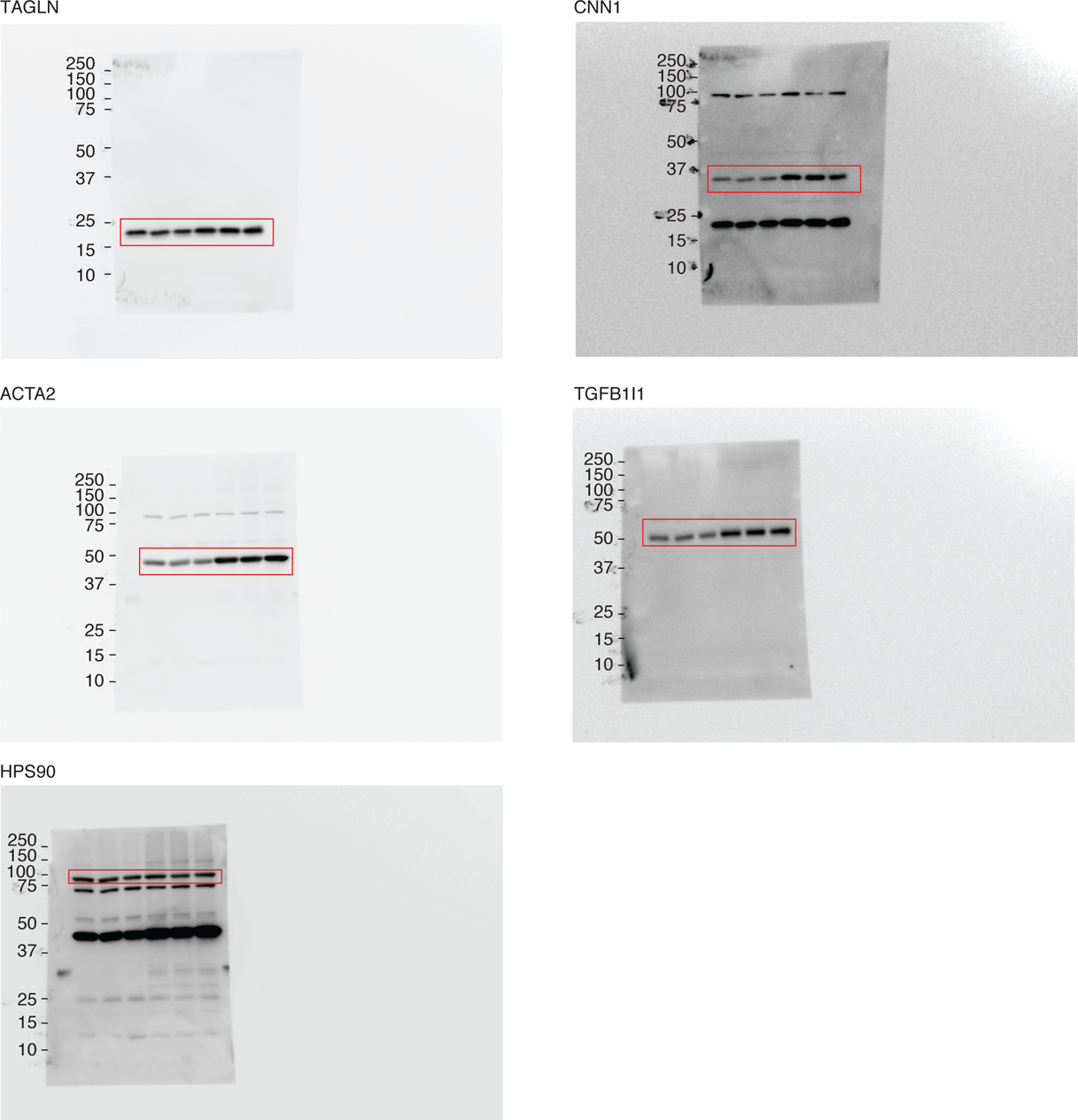

## References

1. Gimona M, Herzog M, Vandekerckhove J, and Small JV. Smooth muscle specific expression of calponin. FEBS Lett. 1990;274(1-2):159–62.

2. Wang X, Hu G, Betts C, Harmon EY, Keller RS, Van De Water L, et al. Transforming growth factor-beta1-induced transcript 1 protein, a novel marker for smooth muscle contractile phenotype, is regulated by serum response factor/myocardin protein. J Biol Chem. 2011;286(48):41589–99.

3. Solway J, Seltzer J, Samaha FF, Kim S, Alger LE, Niu Q, et al. Structure and expression of a smooth muscle cell-specific gene, SM22 alpha. J Biol Chem. 1995;270(22):13460–9.

4. Nanda V, and Miano JM. Leiomodin 1, a new serum response factor-dependent target gene expressed preferentially in differentiated smooth muscle cells. J Biol Chem. 2012;287(4):2459–67.

5. Sawtell NM, and Lessard JL. Cellular distribution of smooth muscle actins during mammalian embryogenesis: expression of the alpha-vascular but not the gamma-enteric isoform in differentiating striated myocytes. J Cell Biol. 1989;109(6 Pt 1):2929–37.

6. Blue EK, Goeckeler ZM, Jin Y, Hou L, Dixon SA, Herring BP, et al. 220- and 130-kDa MLCKs have distinct tissue distributions and intracellular localization patterns. Am J Physiol Cell Physiol. 2002;282(3):C451–60.

7. Kuro-o M, Nagai R, Tsuchimochi H, Katoh H, Yazaki Y, Ohkubo A, et al. Developmentally regulated expression of vascular smooth muscle myosin heavy chain isoforms. J Biol Chem. 1989;264(31):18272–5.

8. Owens GK, Kumar MS, and Wamhoff BR. Molecular regulation of vascular smooth muscle cell differentiation in development and disease. Physiol Rev. 2004;84(3):767–801.

9. Fatica A, and Bozzoni I. Long non-coding RNAs: new players in cell differentiation and development. Nat Rev Genet. 2014;15(1):7–21.

10. Freedman JE, Miano JM, National Heart L, and Blood Institute Workshop P. Challenges and Opportunities in Linking Long Noncoding RNAs to Cardiovascular, Lung, and Blood Diseases. Arterioscler Thromb Vasc Biol. 2017;37(1):21–5.

11. Leung A, Stapleton K, and Natarajan R. Functional Long Non-coding RNAs in Vascular Smooth Muscle Cells. Curr Top Microbiol Immunol. 2016;394:127–41.

12. Miano JM, and Long X. The short and long of noncoding sequences in the control of vascular cell phenotypes. Cell Mol Life Sci. 2015;72(18):3457–88.

13. Ahmed ASI, Dong K, Liu J, Wen T, Yu L, Xu F, et al. Long noncoding RNA NEAT1 (nuclear paraspeckle assembly transcript 1) is critical for phenotypic switching of vascular smooth muscle cells. Proc Natl Acad Sci U S A. 2018;115(37):E8660–E7.

14. Gawad C, Koh W, and Quake SR. Single-cell genome sequencing: current state of the science. Nat Rev Genet. 2016;17(3):175–88.

15. Ounzain S, Micheletti R, Arnan C, Plaisance I, Cecchi D, Schroen B, et al. CARMEN, a human super enhancer-associated long noncoding RNA controlling cardiac specification, differentiation and homeostasis. J Mol Cell Cardiol. 2015;89(Pt A):98–112.

16. Plaisance I, Perruchoud S, Fernandez-Tenorio M, Gonzales C, Ounzain S, Ruchat P, et al. Cardiomyocyte Lineage Specification in Adult Human Cardiac Precursor Cells Via Modulation of Enhancer-Associated Long Noncoding RNA Expression. JACC Basic Transl Sci. 2016;1(6):472–93.

17. Xin M, Small EM, Sutherland LB, Qi X, McAnally J, Plato CF, et al. MicroRNAs miR-143 and miR-145 modulate cytoskeletal dynamics and responsiveness of smooth muscle cells to injury. Genes Dev. 2009;23(18):2166–78.

18. Wang X, Hu G, and Zhou J. Repression of versican expression by microRNA-143. J Biol Chem. 2010;285(30):23241–50.

19. Cheng Y, Liu X, Yang J, Lin Y, Xu DZ, Lu Q, et al. MicroRNA-145, a novel smooth muscle cell phenotypic marker and modulator, controls vascular neointimal lesion formation. Circ Res. 2009;105(2):158–66.

20. Boettger T, Beetz N, Kostin S, Schneider J, Kruger M, Hein L, et al. Acquisition of the contractile phenotype by murine arterial smooth muscle cells depends on the Mir143/145 gene cluster. The Journal of clinical investigation. 2009;119(9):2634–47.

21. Zhou VW, Goren A, and Bernstein BE. Charting histone modifications and the functional organization of mammalian genomes. Nat Rev Genet. 2011;12(1):7–18.

22. He C, Li Z, Chen P, Huang H, Hurst LD, and Chen J. Young intragenic miRNAs are less coexpressed with host genes than old ones: implications of miRNA-host gene coevolution. Nucleic Acids Res. 2012;40(9):4002–12.

23. Wirka RC, Wagh D, Paik DT, Pjanic M, Nguyen T, Miller CL, et al. Atheroprotective roles of smooth muscle cell phenotypic modulation and the TCF21 disease gene as revealed by single-cell analysis. Nature medicine. 2019;25(8):1280–9.

24. Armulik A, Genove G, and Betsholtz C. Pericytes: developmental, physiological, and pathological perspectives, problems, and promises. Developmental cell. 2011;21(2):193–215.

25. Lee G, and Saito I. Role of nucleotide sequences of loxP spacer region in Cre-mediated recombination. Gene. 1998;216(1):55–65.

26. Sun Q, Hao Q, and Prasanth KV. Nuclear Long Noncoding RNAs: Key Regulators of Gene Expression. Trends Genet. 2018;34(2):142–57.

27. Sata M, Maejima Y, Adachi F, Fukino K, Saiura A, Sugiura S, et al. A mouse model of vascular injury that induces rapid onset of medial cell apoptosis followed by reproducible neointimal hyperplasia. J Mol Cell Cardiol. 2000;32(11):2097–104.

28. Kumar A, and Lindner V. Remodeling with neointima formation in the mouse carotid artery after cessation of blood flow. Arterioscler Thromb Vasc Biol. 1997;17(10):2238–44.

29. Clowes AW, Reidy MA, and Clowes MM. Mechanisms of stenosis after arterial injury. Lab Invest. 1983;49(2):208–15.

30. Ballantyne MD, Pinel K, Dakin R, Vesey AT, Diver L, Mackenzie R, et al. Smooth Muscle Enriched Long Noncoding RNA (SMILR) Regulates Cell Proliferation. Circulation. 2016;133(21):2050–65.

31. Holycross BJ, Blank RS, Thompson MM, Peach MJ, and Owens GK. Platelet-derived growth factor-BB-induced suppression of smooth muscle cell differentiation. Circ Res. 1992;71(6):1525–32.

32. Debouck C, and Goodfellow PN. DNA microarrays in drug discovery and development. Nat Genet. 1999;21(1 Suppl):48–50.

## Supplemental References

1. Maass PG, Glazar P, Memczak S, Dittmar G, Hollfinger I, Schreyer L, et al. A map of human circular RNAs in clinically relevant tissues. J Mol Med (Berl). 2017;95(11):1179–89.

2. Hall IF, Climent M, Quintavalle M, Farina FM, Schorn T, Zani S, et al. Circ_Lrp6, a Circular RNA Enriched in Vascular Smooth Muscle Cells, Acts as a Sponge Regulating miRNA-145 Function. Circ Res. 2019;124(4):498–510.

3. You X, Vlatkovic I, Babic A, Will T, Epstein I, Tushev G, et al. Neural circular RNAs are derived from synaptic genes and regulated by development and plasticity. Nat Neurosci. 2015;18(4):603–10.

4. Bolger AM, Lohse M, and Usadel B. Trimmomatic: a flexible trimmer for Illumina sequence data. Bioinformatics. 2014;30(15):2114–20.

5. Kim D, Pertea G, Trapnell C, Pimentel H, Kelley R, and Salzberg SL. TopHat2: accurate alignment of transcriptomes in the presence of insertions, deletions and gene fusions. Genome Biol. 2013;14(4):R36.

6. Liao Y, Smyth GK, and Shi W. featureCounts: an efficient general purpose program for assigning sequence reads to genomic features. Bioinformatics. 2014;30(7):923–30.

7. Mortazavi A, Williams BA, McCue K, Schaeffer L, and Wold B. Mapping and quantifying mammalian transcriptomes by RNA-Seq. Nat Methods. 2008;5(7):621–8.

8. Hong EL, Sloan CA, Chan ET, Davidson JM, Malladi VS, Strattan JS, et al. Principles of metadata organization at the ENCODE data coordination center. Database (Oxford). 2016;2016.

9. Wirka RC, Wagh D, Paik DT, Pjanic M, Nguyen T, Miller CL, et al. Atheroprotective roles of smooth muscle cell phenotypic modulation and the TCF21 disease gene as revealed by single-cell analysis. Nat Med. 2019;25(8):1280–9.

10. Butler A, Hoffman P, Smibert P, Papalexi E, and Satija R. Integrating single-cell transcriptomic data across different conditions, technologies, and species. Nat Biotechnol. 2018;36(5):411–20.

11. Kalluri AS, Vellarikkal SK, Edelman ER, Nguyen L, Subramanian A, Ellinor PT, et al. Single-Cell Analysis of the Normal Mouse Aorta Reveals Functionally Distinct Endothelial Cell Populations. Circulation. 2019;140(2):147–63.

12. Dobin A, Davis CA, Schlesinger F, Drenkow J, Zaleski C, Jha S, et al. STAR: ultrafast universal RNA-seq aligner. Bioinformatics. 2013;29(1):15–21.

13. Li G, Tian L, Goodyer W, Kort EJ, Buikema JW, Xu A, et al. Single cell expression analysis reveals anatomical and cell cycle-dependent transcriptional shifts during heart development. Development. 2019;146(12).

14. Kannan S, Miyamoto M, Lin BL, Zhu R, Murphy S, Kass DA, et al. Large Particle Fluorescence-Activated Cell Sorting Enables High-Quality Single-Cell RNA Sequencing and Functional Analysis of Adult Cardiomyocytes. Circ Res. 2019;125(5):567–9.

15. Wang L, Yu P, Zhou B, Song J, Li Z, Zhang M, et al. Single-cell reconstruction of the adult human heart during heart failure and recovery reveals the cellular landscape underlying cardiac function. Nat Cell Biol. 2020;22(1):108–19.

16. Skelly DA, Squiers GT, McLellan MA, Bolisetty MT, Robson P, Rosenthal NA, et al. Single-Cell Transcriptional Profiling Reveals Cellular Diversity and Intercommunication in the Mouse Heart. Cell Rep. 2018;22(3):600–10.

17. Asp M, Giacomello S, Larsson L, Wu C, Furth D, Qian X, et al. A Spatiotemporal Organ-Wide Gene Expression and Cell Atlas of the Developing Human Heart. Cell. 2019;179(7):1647–60 e19.

18. Thorvaldsdottir H, Robinson JT, and Mesirov JP. Integrative Genomics Viewer (IGV): high-performance genomics data visualization and exploration. Brief Bioinform. 2013;14(2):178–92.

19. Ayari H, and Bricca G. Identification of two genes potentially associated in iron-heme homeostasis in human carotid plaque using microarray analysis. J Biosci. 2013;38(2):311–5.

20. Bekelis K, Kerley-Hamilton JS, Teegarden A, Tomlinson CR, Kuintzle R, Simmons N, et al. MicroRNA and gene expression changes in unruptured human cerebral aneurysms. J Neurosurg. 2016;125(6):1390–9.

21. Ballantyne MD, Pinel K, Dakin R, Vesey AT, Diver L, Mackenzie R, et al. Smooth Muscle Enriched Long Noncoding RNA (SMILR) Regulates Cell Proliferation. Circulation. 2016;133(21):2050–65.

22. Trapnell C, Roberts A, Goff L, Pertea G, Kim D, Kelley DR, et al. Differential gene and transcript expression analysis of RNA-seq experiments with TopHat and Cufflinks. Nat Protoc. 2012;7(3):562–78.

23. Lyu Q, Xu S, Lyu Y, Choi M, Christie CK, Slivano OJ, et al. SENCR stabilizes vascular endothelial cell adherens junctions through interaction with CKAP4. Proc Natl Acad Sci U S A. 2019;116(2):546–55.

24. Ahmed ASI, Dong K, Liu J, Wen T, Yu L, Xu F, et al. Long noncoding RNA NEAT1 (nuclear paraspeckle assembly transcript 1) is critical for phenotypic switching of vascular smooth muscle cells. Proc Natl Acad Sci U S A. 2018;115(37):E8660–E7.

25. Wang X, Hu G, Gao X, Wang Y, Zhang W, Harmon EY, et al. The induction of yes-associated protein expression after arterial injury is crucial for smooth muscle phenotypic modulation and neointima formation. Arterioscler Thromb Vasc Biol. 2012;32(11):2662–9.

26. Wang X, Hu G, Betts C, Harmon EY, Keller RS, Van De Water L, et al. Transforming growth factor-beta1-induced transcript 1 protein, a novel marker for smooth muscle contractile phenotype, is regulated by serum response factor/myocardin protein. J Biol Chem. 2011;286(48):41589–99.

27. Liu F, Wang X, Hu G, Wang Y, and Zhou J. The transcription factor TEAD1 represses smooth muscle-specific gene expression by abolishing myocardin function. J Biol Chem. 2014;289(6):3308–16.

28. Wang X, Hu G, and Zhou J. Repression of versican expression by microRNA-143. J Biol Chem. 2010;285(30):23241–50.

29. Xu F, Ahmed AS, Kang X, Hu G, Liu F, Zhang W, et al. MicroRNA-15b/16 Attenuates Vascular Neointima Formation by Promoting the Contractile Phenotype of Vascular Smooth Muscle Through Targeting YAP. Arterioscler Thromb Vasc Biol. 2015;35(10):2145–52.

30. Osman I, He X, Liu J, Dong K, Wen T, Zhang F, et al. TEAD1 (TEA Domain Transcription Factor 1) Promotes Smooth Muscle Cell Proliferation Through Upregulating SLC1A5 (Solute Carrier Family 1 Member 5)-Mediated Glutamine Uptake. Circ Res. 2019;124(9):1309–22.

